# A complete suite for compact programming of protein secretion

**DOI:** 10.1101/2023.10.04.560774

**Authors:** Alexander E. Vlahos, Connor C. Call, Noah Eckman, Samarth E. Kadaba, Siqi Guo, Abby R. Thurm, Jeewoo Kang, Eric A. Appel, Lacramiora Bintu, Xiaojing J. Gao

## Abstract

Synthetic biology has developed powerful tools to program complex behaviors, often using genetic control. Protein circuits offer a compact alternative, yet applications with intercellular signals often lack key regulatory capabilities and tunability. Here, we employ a parts-based engineering strategy to develop a single processing and output module for synthetic protein circuits, enabling complex logic, tunable sensitivity, and control over output magnitude. Using high-throughput assays, we systematically analyze the impact of human transmembrane domains on surface expression and circuit performance. We demonstrate the utility of these optimizations by encoding an open-loop circuit within translational delivery vectors, including viral and mRNA platforms, and validate its performance in vivo. Furthermore, we demonstrate multi-input logic and showcase a novel, protein-level NIMPLY gate to regulate CAR T-cell activation. Our modular design strategy provides new insights into domain-based protein engineering and establishes a versatile and complete protein-level platform to control intercellular signaling for translational cell therapies.

**Graphical Abstract:** 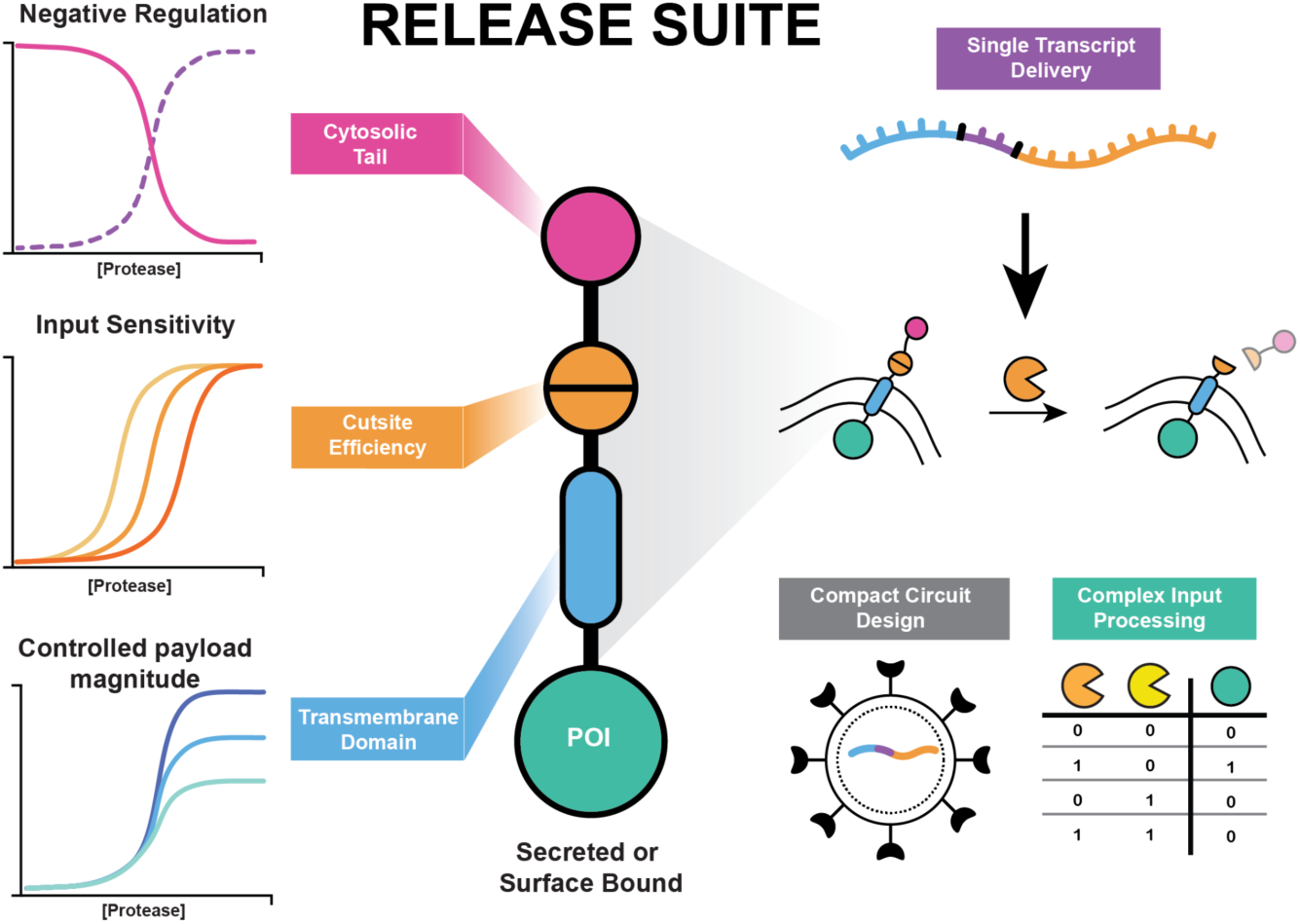

## Introduction

The field of synthetic biology applies engineering principles to program cellular behavior by constructing biomolecular circuits that mimic natural regulatory systems^1–5^. These circuits impart cells with new behaviors such as the ability to sense specific ligands in the microenvironment^6–9^, process this information, and execute therapeutic functions^10,11^ in a predictable and controllable manner. Historically, synthetic gene circuits had dominated the field^11–15^, built from extensive promoter engineering^16,17^, the optimization of synthetic transcription factors^18^, and other regulatory DNA binding elements^19^ to modulate gene expression with high specificity and programmability. More recently, synthetic protein circuits have emerged as a powerful alternative, leveraging orthogonal proteases to implement signal-processing capabilities including Boolean logic operation and quantitative processing, entirely at the protein level^20,21^. To engage intercellular communications, protein secretion circuits were then further developed that use cytosolic proteases to control extracellular payloads’ transit through the secretory compartments^22–25^. Protein circuits offer distinct advantages over traditional gene circuits, such as faster responsive kinetics, reduced genetic payloads, and improved compatibility with translational delivery platforms like mRNA and viral vectors^26,27^.

While the existing protease circuit platforms are nominally “complete” for programming protein secretion, i.e., capable of performing all possible quantitative tunings and logic operations, there is a major tension between completeness and the practical value of such circuits. This tension is compounded by most *in vivo* demonstrations relying on DNA transient transfections^24,25,28^, underscoring an unmet need to address delivery constraints. In the existing platforms, even a simple task such as flipping the sign or tuning the threshold of a regulation requires additional orthogonal proteases^20,29^. Such additions compromise circuit compactness that is essential for their therapeutic deliveries; they also create layered architecture through which circuit performance tends to deteriorate. Therefore, there remained a critical gap with regard to maintaining the advantages of protease circuits while enabling functionalities that actually rival those of gene circuits.

To address these limitations, we focused on expanding RELEASE (Retained Endoplasmic Cleavable Secretion), a previously described platform that enables protease-dependent control of protein secretion and membrane display through the proteolytic removal of cytosolic ER-retention motifs^22^ (**Fig. 1a, b**). While other platforms have utilized cytosolic ER-retention mechanisms to regulate secretion^23–25^, RELEASE offered initial examples of compact implementation of logic operation by integrating them into the output module^22^. We hypothesized that RELEASE could serve as an ideal foundation for creating more sophisticated and compact protein circuits if its core components were systematically engineered.

**Figure 1:**
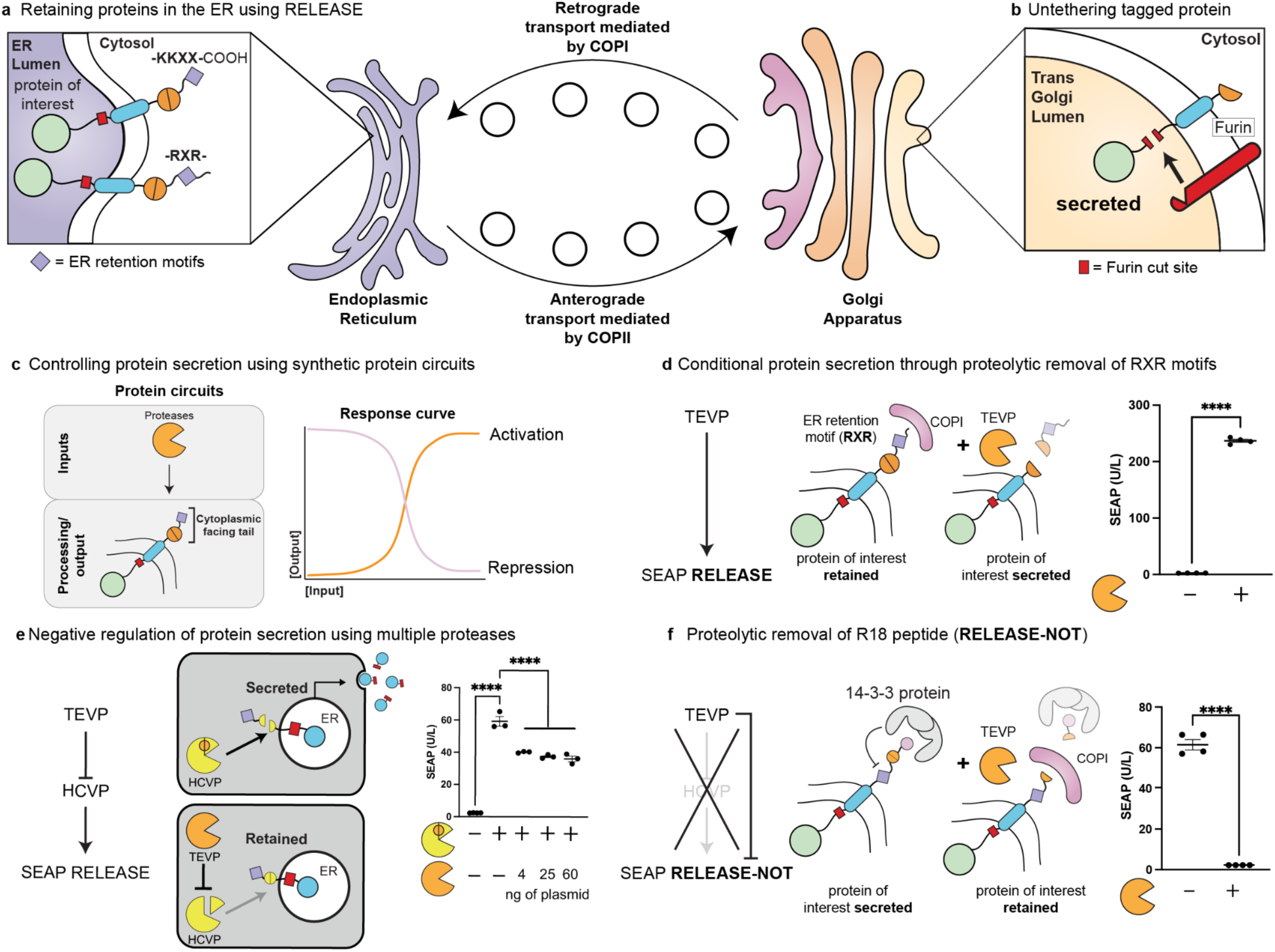
Repression of protein secretion using 14-3-3 proteins. **a)** Schematic of the **<u>R</u>**etained **<u>E</u>**ndoplasmic C**<u>lea</u>**vable **<u>Se</u>**cretion (RELEASE) platform using native ER-retention motifs (purple diamond) retain tagged proteins of interest in the ER. Only through the activation or expression of a protease such as TEVP (orange pac-man) will there be proteolytic removal of the ER-retention motif, and subsequent transport through the conventional secretory pathway. **b)** Upon reaching the trans-golgi apparatus, tagged proteins of interest are untethered from the membrane via proteolytic cleavage by the lumenal-facing native furin endoprotease. **c**) Schematic highlighting the activation or repression of proteins of interest by modifying the cytoplasmic tail of RELEASE. **d)** Additional ER-retention motifs such as the diarginine (-RXR-) motif are compatible with the RELEASE platform. Co-expression of proteases such as TEVP significantly increased SEAP secretion. **e**) Using multiple proteases to repress protein secretion in response to the activation of an input protease. The intermediate protease (HCVP - yellow pac-man) is inactivated by the expression of the input protease (TEVP), and increasing the amount of TEVP significantly reduced SEAP secretion, but could not reach basal levels. **f**) In contrast, through harnessing the proteolytic removal of the R18 peptide (pink circle), negative regulation of protein secretion could be achieved using a single input protease. When TEVP is absent, the R18 peptide recruits endogenous 14-3-3 scaffolding proteins to block the interaction between RELEASE and the COPI retrograde complex, resulting in the payload being secreted. When TEVP is present, the R18 peptide is removed and the retention properties of RELEASE are restored. Each dot represents a biological replicate. Mean values were calculated from four replicates. The error bars represent +/- SEM. The results are representative of at least two independent experiments; significance was tested using an unpaired two-tailed Student’s *t*-test between the two indicated conditions for each experiment. For experiments with multiple conditions, a one-way ANOVA with a Tukey’s post-hoc comparison test was used to assess significance. \*\*\*\**p* < 0.0001.

Inspired by part-optimization strategies used to enhance other synthetic biology platforms such as SNIPR^30^ and MESA^31^, we adopted a modular, parts-based approach to uncover the design principles of RELEASE. For example, by expanding the types and arrangement of cytosolic-facing domains, we envision enabling more complex localization control through the coordinated recruitment of endogenous scaffolding proteins^32,33^, mimicking natural post-translational mechanisms. Additionally, key components such as the transmembrane domain, which may significantly affect protein expression^34^, trafficking^35^, and function remain largely unexplored in the context of synthetic expression platforms, such as RELEASE. By systematically optimizing these modular components, we aimed to establish a more compact, tunable, and generalizable platform for programming protein secretion, with capabilities analogous to synthetic gene circuits while maintaining or even improving compatibility with translational delivery vectors.

Here, we introduce a complete suite of RELEASE variants and design principles to program protein secretion. Inspired by the capabilities of synthetic gene circuits, we demonstrate negative regulation of protein secretion, quantitative input processing, expression level tuning, and functional completeness of multi-input logic gating, entirely at the protein level. We also focus on the efficient delivery of synthetic protein circuits, using polycistronic designs compatible with viral vectors and mRNA delivery to engineer cells with inducible control over protein secretion both in vitro and in vivo. Finally, we apply these designs to achieve multi-input control of CAR surface expression and activation, highlighting potential for enabling conditional, logic-based control of immunotherapies. The RELEASE suite is a compact and modular protein-level processing and output platform that expands the synthetic protein circuit toolkit and enables circuit-level control of protein secretion while minimizing the overall genetic payload.

## Results

### Proteolytic repression of protein secretion enabled by 14-3-3 proteins

To control protein secretion at the genetic level, synthetic genetic circuits can tune gene expression using transcription factors as direct inputs, which can act as activators or repressors. Unlike genetic circuits that regulate transcription, RELEASE controls secretion through proteolytic removal of ER-retention motifs (**Fig. 1c**). However, these current tools can only activate protein secretion in response to protease activity and not repress it (i.e. NOT logic). Furthermore, the original RELEASE platform used a cytoplasmic dilysine ER-retention motif (-KKXX-CO_2_H)^36^ that only functioned when placed at the C-terminus^22^. This strict spatial requirement limits RELEASE from being compatible with additional cytoplasmic-facing functional protein motifs^37^, or with self-cleaving 2A tags in polycistronic genes^38^. For example, some membrane proteins control their surface expression by regulating interactions between ER-retention motifs and the COPI coatomer through the recruitment of endogenous 14-3-3 proteins^32,33,39,40^ (**Supplementary Fig 1**). Modifying the RELEASE platform to enable functionalization of the C-terminus, while preserving the ER-retention capabilities would enhance programmability of protein-level control of protein secretion.

To overcome this constraint, we used an alternative ER-retention motif known as the diarginine motif (-RXR-)^41,42^, which functions identically to the dilysine motif (**Fig. 1b**), but is not restricted to the C-terminus (**Supplementary Fig 2**). Using these new RELEASE constructs, we transiently transfected HEK293 cells and validated the direct activation of Secreted Alkaline Embryonic Phosphatase (SEAP) secretion when co-expressing the cognate protease (i.e. TEVP) relative to its absence (**Fig. 1d**). As previously shown^22^, the modular nature of RELEASE enables additional variants to be created by switching the cytosolic protease cut site to make a hepatitis C viral protease (HCVP)-responsive RELEASE (**Supplementary Fig 3**). For the remainder of this study, we used -RXR-based RELEASE variants.

Since RELEASE can only be used to increase protein secretion in response to protease activity, we first attempted to enable NOT logic using a two-protease circuit^20,29^ (**Fig 1e** - left panel). This circuit used TEVP as the input protease, which inhibited an intermediate protease (HCVP - yellow pacman) from activating the cognate RELEASE construct. By titrating the amount of TEVP, we observed a significant and dose-dependent decrease in SEAP secretion; however, even at high levels of TEVP, SEAP secretion was not entirely repressed (**Fig. 1e**). Rather than optimizing the two-protease circuit for NOT logic, we sought to directly program this behavior into RELEASE by leveraging endogenous 14-3-3 protein recruitment to modulate ER-retention activity (**Supplementary Fig. 1**). Previously, the R18 peptide has been described to bind the conserved amphipathic ligand groove of 14-3-3 proteins^43^ via the WLDLE motif^44^. We hypothesized that fusing R18 to the C-terminus of RELEASE would sufficiently block ER-retention and drive protein secretion. Indeed, the addition of R18 effectively blocked the retention capabilities of RELEASE and resulted in a significant increase in SEAP secretion relative to the RELEASE construct with a R18 mutant peptide incapable of recruiting 14-3-3 (**Supplementary Fig 4**).

Following this validation, we placed a protease cut site (such as TEVP) between the ER-retention motif and R18 peptide to create RELEASE-NOT (**Fig. 1f**). RELEASE-NOT relies on the recruitment of 14-3-3 proteins to enable constitutive secretion of a payload, until proteolytic removal of the R18 peptide results in restoration of ER retention activity (**Fig. 1f**). Co-expression of TEVP with a TEVP-responsive RELEASE-NOT significantly reduced protein secretion relative to when the protease was absent (**Fig. 1f**), more streamlined than the double-negative configuration (**Fig. 1e**) while substantially outperforming the latter. Like RELEASE, a HCVP-responsive RELEASE-NOT was engineered, by switching the protease cut site (**Supplementary Fig. 5**). We also tested additional phosphorylation-dependent motifs to recruit endogenous 14-3-3 proteins^37^, which showed comparable activity to the R18 peptide (**Supplementary Fig. 6**). However, for the rest of this work, we exclusively used the R18 peptide because it was not constrained to the C-terminus^37^, and its phosphorylation-independent activity was hypothesized to make it less sensitive to changes in cell types or states. These results show the potential of incorporating additional protein motifs at the C-terminus of RELEASE to impart new processing capabilities.

### Compact quantitative processing of inputs

Considering that some intercellular signals such as cytokines and growth factors have pleiotropic activity based on their concentrations^45–47^, the ability to tune the sensitivity of circuit activation may be an important capability for immune regulation or tissue regeneration^47^. Synthetic gene circuits can accomplish this by changing the interaction between synthetic transcription factors^18,48^ and their target sites to tune the response threshold of a given gene of interest. For example, modifying DNA-binding domains^49^ or introducing cooperative binding sites^50^ can fine-tune activation sensitivity. Similarly, at the protein level, multiple orthogonal proteases can be engineered to quantitatively tune protein expression within defined input regions to achieve bandpass behaviour^20^. However, if we could predictably tune the input sensitivity of RELEASE by manipulating protease cut site efficiency (**Fig. 2a**), we may be able to achieve the same quantitative behaviors without the need for additional proteases^20^. This would effectively streamline the entire circuit, reducing the overall genetic payload.

**Figure 2:**
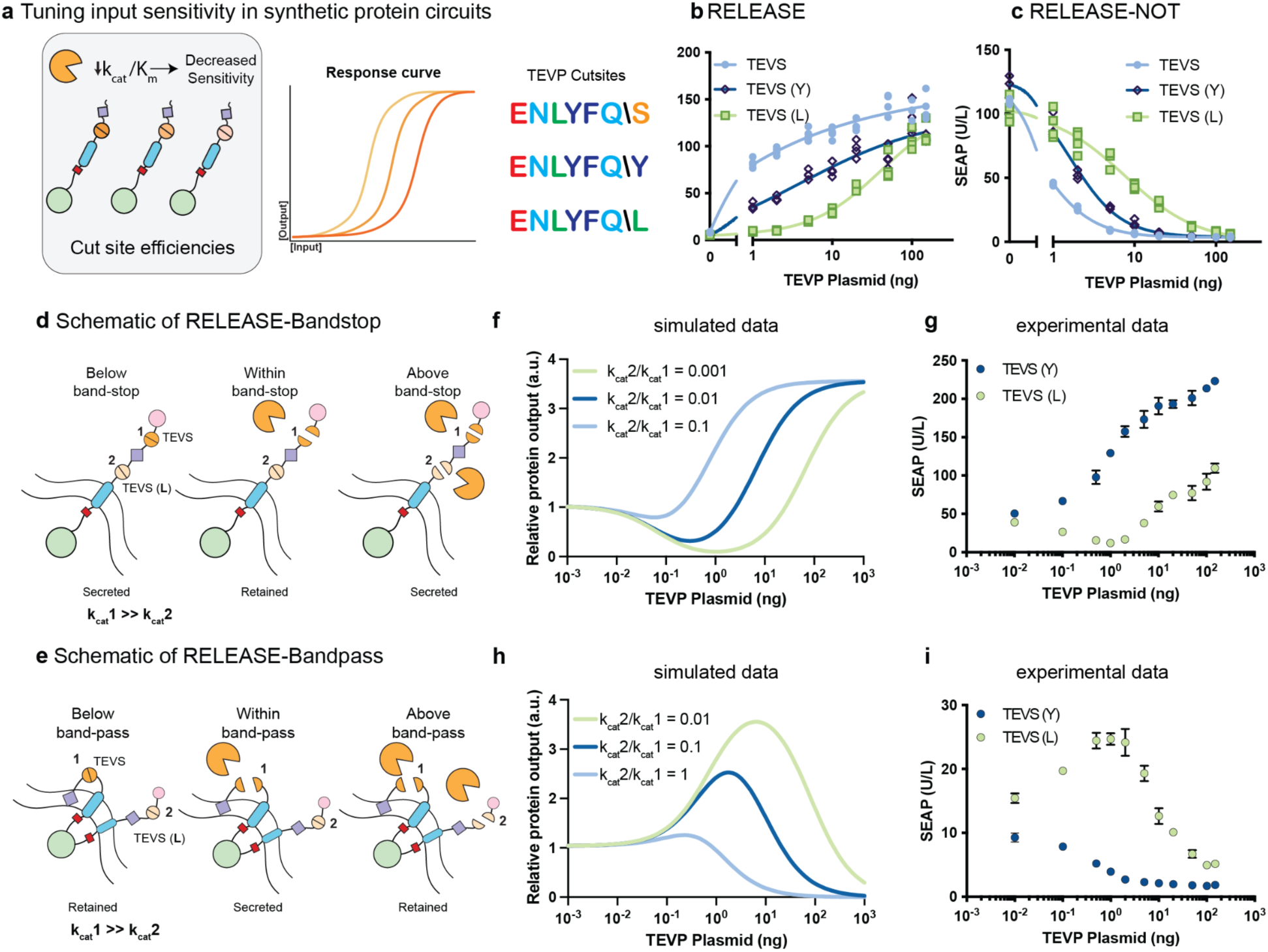
Compact quantitative processing of inputs. **a)** Schematic of tuning input sensitivity in synthetic protein circuits. Canonical cut site of TEVP is ENLYFQ\S, where S is the P1’ position that can be mutated to change cleavage efficiency. **b)** Input-output curves of three RELEASE constructs with modified amino acids at the P1’ position titrated with increasing amounts of the TEVP plasmid. Changing the canonical sequence reduced the cleavage activation and the activation threshold required for protein secretion. **c**) RELEASE-NOT observed similar trends when mutating the P1’ position of the TEVP cut site. The data is fitted to a Hill equation with a variable slope represented by the respective lines, and the EC_50_s and IC_50_s for each variant can be found in **Supplementary Table 1**. Schematics outlining the different states of **d)** RELEASE-Bandstop and **e)** RELEASE-Bandpass, highlighting if the tagged protein payload would be retained or secreted. The purple diamond represent ER retention motifs and the pink circle represents the R18 peptide. **f)** Simulated data of input-output curves for the RELEASE-Bandstop variant, simulating differences in outputs by changing the ratio between the cleavage efficiencies between the two cut sites. **g)** Input-output curve of RELEASE-bandstop titrated with TEVP plasmid. **h)** Simulated data of input-output curves for the RELEASE-Bandpass variant, simulating differences in outputs by changing the ratio between the cleavage efficiencies between the two cut sites. **i)** Input-output curve of RELEASE-bandpass titrated with TEVP plasmid. Each dot represents a biological replicate (**b and c**). Mean values were calculated from 4 replicates (**b,c, g, and i**). The error bars represent +/- SEM. The results are representative of at least two independent experiments.

Previously, the cleavage efficiency of RELEASE was manipulated by altering the number of transmembrane domains or amino acid residues flanking the protease cut site^22^. However, neither strategy yielded a consistent or predictable framework for tuning secretion thresholds. To simplify the design process, we focused on modulating cleavage efficiency through changes to the canonical amino acid sequence of the protease cut site. We selected the TEVP-inducible RELEASE construct as our test case because the P1’ position of the canonical cut site (e.g. ENLYFQ/S) has been well documented to influence cleavage efficiency^51,52^. Using three different cut site variants, we validated that we could shift the activation of RELEASE (**Fig. 2b**) and RELEASE-NOT (**Fig. 2c**). The shift in activation or repression using RELEASE, or RELEASE-NOT aligned with the published trends^51^, where the canonical site was the most efficiently cleaved, followed by tevs(Y), and finally tevs(L) (**Supplementary Table 1**).

Having established predictable control over the cleavage efficiencies of RELEASE and RELEASE-NOT, we next aimed to achieve more complex analog signal processing such as bandstop and bandpass control of protein secretion using a single protease input. We integrated protein components designed to activate or repress protein secretion into two RELEASE variants, which each contained two protease cut sites that could be cleaved by the same protease, but with different cleavage efficiencies (**Fig. 2d,e**). RELEASE-bandstop combined the c-terminal tails of RELEASE and RELEASE-NOT and was designed to drive protein secretion at low and high protease concentrations, while intermediate levels of protease activity led to protein retention (**Fig. 2d**). We computationally simulated the input-output curves of this circuit given different cleavage efficiency ratios between the two cut sites (**Fig. 2f**), and validated the trends in experiments, where we titrated the amount of TEVP co-transfected with a constant level of the bandstop reporter in HEK293 cells (**Fig. 2g**).

To engineer RELEASE-bandpass, we replaced the C-terminus of the previously described AND-gate RELEASE^22^ construct with RELEASE-NOT (**Fig. 2e**). Since the N-terminal ER retention motif is directly anchored to the ER membrane^53,54^, and not COPI-mediated, its retention capabilities was unaffected by the recruitment of 14-3-3 proteins via the R18 peptide. This meant that under low and high protease concentrations the tagged payload would be retained, and only under intermediate concentrations would we get secretion. We implemented a similar strategy to RELEASE-bandstop, where we first modelled different cleavage efficiency ratios (**Fig. 2h**) to identify those exhibiting bandpass behavior, and then experimentally validated the design of RELEASE-bandpass (**Fig. 2i**). This work highlights proof-of-principle quantitative processing using RELEASE variants to control protein secretion without the need of additional proteases.

### High-throughput transmembrane domain characterization to modulate payload magnitude

Precise control over input responsiveness is critical, but equally important is the ability to regulate the magnitude of the secreted output. Synthetic gene circuits can tune the expression of their outputs by using promoters of different strengths. Achieving the same level of control at the post-translational level is more challenging, as it often requires engineering protein stability via incorporating degrons^55,56^ or manipulating post-translational modifications^57^, which may affect protein function. To address this challenge without introducing additional functional domains, we sought to regulate protein secretion by modifying the transmembrane domain (TMD) of RELEASE (**Fig. 3a**). Given its critical role in ER membrane insertion^58^, the transport and sorting of integral membrane proteins^34,35^, and its effect on the surface expression of synthetic receptors^31,59^, we hypothesized that the TMD could serve as a tunable element for controlling protein secretion with RELEASE.

**Figure 3:**
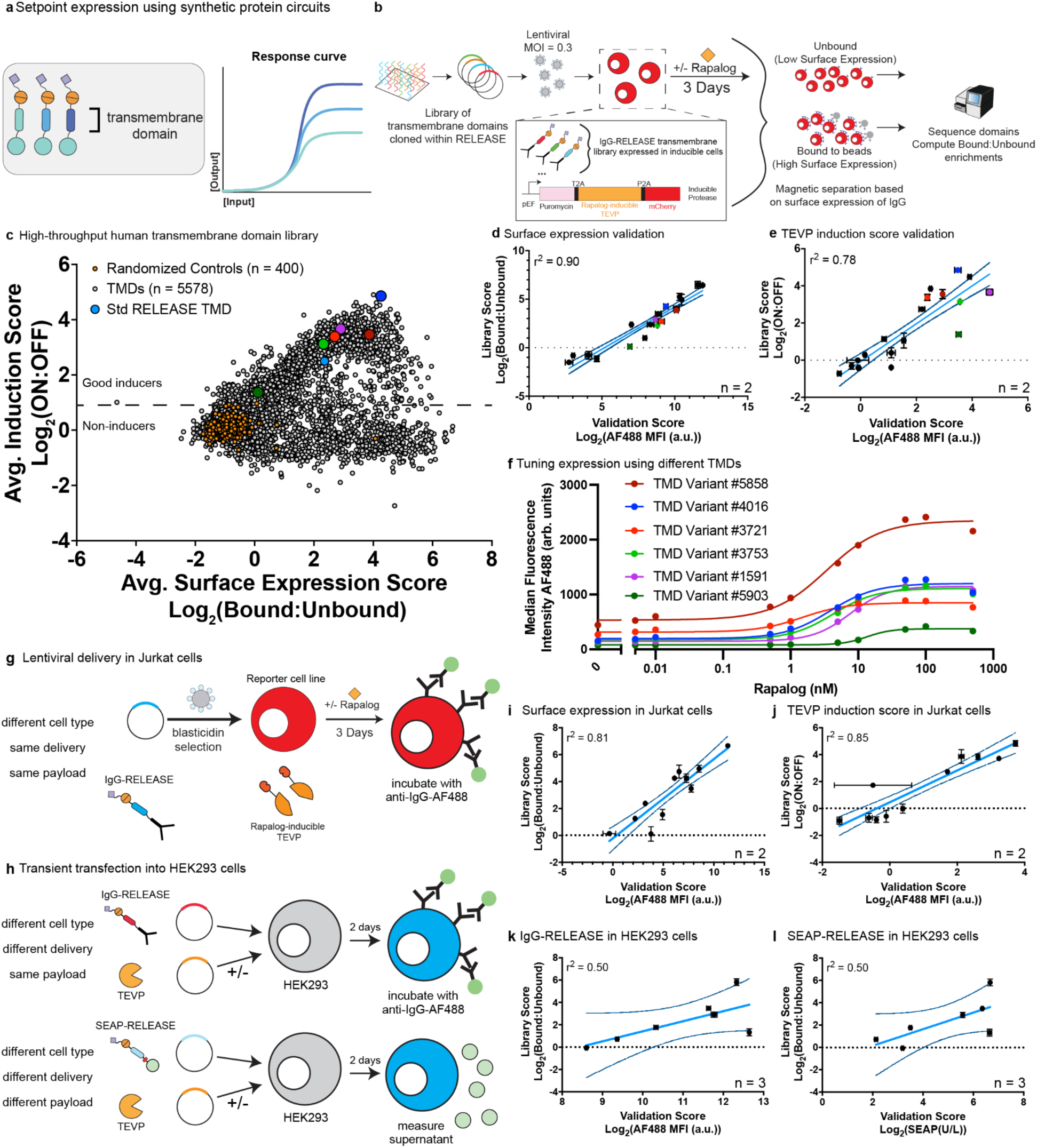
High-throughput human transmembrane characterization modulates protein expression. **a)** Schematic representation of tuning steady-state set points of protein expression using synthetic gene or protein circuits. **b)** Schematic of high-throughput pull down assay. A pooled library of RELEASE constructs with different TMDs was synthesized, cloned and delivered into reporter cells. The reporter cells contain a rapalog-inducible TEVP to induce surface expression of a synthetic surface marker (IgG-RELEASE). Cells were treated with and without rapalog for 3 days before magnetic bead separation, followed by sequencing. **c)** The high-throughput dataset for TMD variants bifurcated when plotting the average TEVP induction scores with the average surface expression scores. Negative controls were composed of randomized amino acid sequences of 21 amino acids. The original RELEASE construct and a small subset used in subsequent individual assays were highlighted. The dotted line represents the threshold enrichment score (>3 standard deviations from the negative population induction score average) to determine TMDs that were good inducers vs. non-inducers. **d)** Correlation of high-throughput measurements for surface expression scores with individual surface staining experiments (r^2^ = 0.90, N = 20 TMDs, n = 2 biological replicates). **e)** Correlation of high-throughput measurement for TEVP induction scores with individual surface staining experiments (r^2^ = 0.78, N = 19 TMDs, n = 2 biological replicates). **f)** Six TMD variants with log_2_[TEVP induction scores] > 2 were used to validate that IgG surface expression could be tuned by different TMDs. EC_50_s for each variant can be found in **Supplementary Table 4**. **g)** Schematic representation of IgG-RELEASE TMD variants delivered into Jurkat cells via lentiviral transduction. **i)** Correlation of high-throughput measurements with individual surface staining experiments for surface expression (r^2^ = 0.81, N = 11 TMDs, n = 2 biological replicates). **j**) Correlation of high-throughput measurements for TEVP induction scores with individual surface staining experiments (r^2^ = 0.85, N = 11 TMDs, n = 2 biological replicates). **h)** Schematic representation of IgG-RELEASE or SEAP-RELEASE delivered into adherent HEK293 cells via transient transfection. **k)** Correlation of high-throughput measurements with individual surface staining measurements for surface expression (r^2^ = 0.50, N = 7 TMDs, n = 3 biological replicates). **l)** Correlation of high-throughput measurements for surface expression with individual SEAP secretion measurements (r^2^ = 0.50, N = 7 TMDs, n = 3 biological replicates). For each correlation, the dotted lines denote 95% confidence intervals.

To screen the effect of TMDs on RELEASE activity, we coupled the synthetic surface marker (FC domain of human IgG1) from the HT-Recruit assay^60^ to the RELEASE platform (IgG-RELEASE, **Supplementary Fig. 7**)). A library of IgG-RELEASE variants was generated using the online UNIPROT database to select for TMDs from human integral membrane proteins that had signal peptides. For multi-spanning transmembrane proteins containing signal peptides, each transmembrane domain was treated as an individual library element. To mirror the naming convention of transmembrane proteins, we use Type I TMDs to refer to those present in their native orientations, and Type II TMDs for those whose intramembrane orientation is flipped to to ensure compatibility with RELEASE (**Supplementary Fig. 7b**). Since the juxtamembrane sequence can affect protein expression^30^ we included two variants for each Type I TMD - one with and one without its native juxtamembrane sequence. For Type II TMDs, we appended the juxtamembrane sequence of CD8ɑ, as the native C-terminal residues immediately after the TMD were not the true juxtamembrane residues (**Supplementary Fig. 7c**). We also included 400 random amino acid sequences to serve as negative controls. The entire library spanned approximately 6000 different IgG-RELEASE variants with different TMDs and a complete list of amino acid sequences used for the library can be found in **Supplementary Table 2**. The library was delivered into a K562 reporter cell line containing a rapalog-inducible TEVP (**Fig. 3b**).

We screened the TMD library for surface expression, ER retention, and TEVP inducibility. The pooled library of cells was separated into two groups: one induced with rapalog, where TEVP was active (ON population) and the other treated with a vehicle control, where TEVP was inactive (OFF population). Cells were incubated for 3 days so that the payload level on the cell surface reached steady-state and then we performed magnetic sorting (**Fig. 3b**). As previously described^60^, the TMD sequences were then analyzed, and the read counts for the bound and unbound fractions were used to compute an enrichment score, defined as Log_2_(Bound/Unbound), for each library element. Over 93% of the original sequences met the count threshold, yielding data for 5,578 distinct TMD variants. The enrichment scores for TMDs in the ON population provided information on the absolute surface expression of IgG (**Supplementary Fig. 8a**), while the fold enrichment in the OFF population was indicative of ER retention capabilities of TMDs used with RELEASE (**Supplementary Fig. 8b**). We then took the ratio of these two enrichment scores to represent TEVP inducibility (**Supplementary Fig. 8c**). Finally, we validated that these measurements showed high reproducibility between biological replicates (**Supplementary Fig. 8**). The choice of TMD substantially affected all three of the aforementioned scores.

To further guide the engineering of RELEASE using different TMDs, we plotted the average scores between surface expression and TEVP inducibility and observed a bifurcation in our data, where some TMDs were classified as good inducers and others as non-inducers (**Fig. 3c**). In general, the presence of juxtamembrane sequences drastically improved TEVP induction (**Supplementary Fig. 9a**), while type I TMDs outperformed type II TMDs (**Supplementary Fig. 9b**). We individually validated a small subset of TMD variants and observed strong correlations with our high-throughput measurements for surface expression (r^2^ = 0.90, **Fig. 3d**), and TEVP induction (r^2^ = 0.78, **Fig. 3e**). Using different TMDs, we confirmed that RELEASE can fine-tune protein surface expression at steady-state with minimal effects on input sensitivity (**Fig. 3f**).

To assess the generalizability of our high-throughput measurements, we tested additional subsets of TMDs in different cell types (i.e. Jurkat cells - **Fig. 3g**, or HEK293 cells - **Fig. 3h**), across different delivery methods (lentiviral transduction vs. transient transfection), and with different payloads (IgG vs. SEAP). In Jurkat cells, we observed strong correlations in surface expression (r^2^ = 0.81, **Fig. 3i**), and TEVP induction (r^2^ = 0.85, **Fig. 3j**). In HEK293 cells, transient transfections yielded moderate correlations with the library data (r^2^ = 0.50, **Fig. 3k**). We also observed a moderate correlation when using a secreted protein payload instead of a surface-expressed one (r^2^ = 0.50, **Fig. 3l**). These general trends also held up with other RELEASE variants, like RELEASE-NOT (**Supplementary Fig. 10**). As expected, while the behavior of the secretory pathway depends on cell types, there seems to be some context-independent commonality as well. Our dataset would help nominate TMD variants for small scale re-screenings when we port RELEASE into other cell types. While this study emphasizes the engineering of RELEASE, further analysis of our dataset may yield broader insights into membrane protein biology.

### Polycistronic cassettes encode protein circuits for viral vectors

As we expand the functionality of RELEASE, we are keenly aware that their delivery remains a key challenge in their translation for cell engineering. Traditionally, to engineer cells, synthetic circuits are delivered via viral vectors such as lentiviruses or adeno-associated viruses (AAV) (**Fig. 4a**). However, a major limitation of viral vectors is their limited genomic cargo capacity^61^, which can hinder the delivery of large or complex synthetic circuits. AAVs, for instance, have a packaging limit of approximately 4.8 kb^62^, while lentiviruses can accommodate around 8–10 kb^63^. Exceeding these limits can compromise vector stability, reduce transduction efficiency, and hinder proper expression of the synthetic circuit^64^. In comparison, the post-translational nature of synthetic protein circuits enables them to be compactly encoded within a single polycistronic gene, under the control of a single promoter where each protein component is separated using self-cleaving 2A peptides^38,65^. This compact design helps overcome the delivery bottleneck; however, precisely controlling the stoichiometry of individual protein components in a single-transcript design remains challenging, as all components are translated together^26^. Additionally, the activity of some proteases, such as HCVP, can be modulated using small molecule inhibitors, such as Asunaprevir (ASV)^66,67^, effectively altering protease activity and concentration.

**Figure 4:**
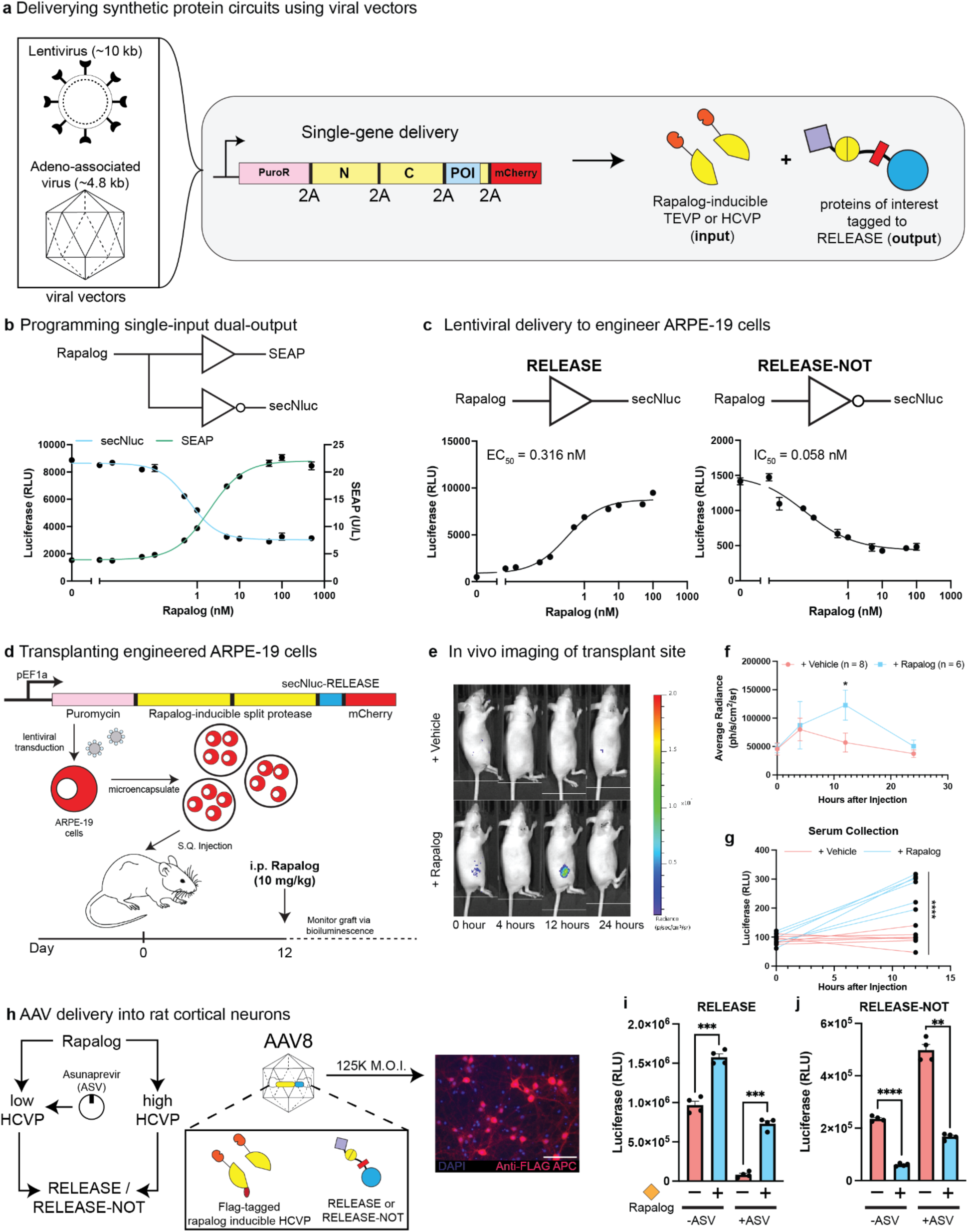
Polycistronic gene cassettes encode protein circuits for cellular engineering. **a)** Schematic of using viral vectors to deliver synthetic protein circuits. **b)** A polycistronic gene encoding a synthetic protein circuit with the antagonistic control of two secreted reporter proteins in response to rapalog that was stably integrated into HEK293 cells via lentiviral transduction. **c)** ARPE-19 cells were stably integrated with a single polycistronic gene encoding a synthetic protein circuit that activated or repressed secNluc secretion when incubated with rapalog, using RELEASE or RELEASE-NOT, respectively. **d)** Schematic of in vivo delivery of engineered ARPE-19 cells. **e)** In vivo imaging of secNluc in animals subcutaneously transplanted with microencapsulated engineered ARPE-19 cells at day 12. **f)** Dose-response curves of average radiance measured using the in vivo imager after injection rapalog or PBS via retro-orbital injection. **g)** Following injection with rapalog or PBS, the serum was collected via the tail-vein and secNluc was measured. Animals injected with rapalog showed a significant increase in the amount of circulating secNluc in the serum compared to animals injected with the PBS control. **h)** Schematic of AAV8 delivery of synthetic protein circuit to control protein secretion into rat cortical neurons. The activity of HCVP was modulated by incubating cells with the Asunaprevir (ASV) inhibitor. In addition, the HCVP used to induce RELEASE contained a FLAG epitope, which enabled us to stain the rat-cortical neurons using an anti-FLAG antibody to confirm the delivery of the synthetic protein circuit. Scale bar = 100 µM. **i)** Secreted luciferase was measured in rat cortical neurons engineered with synthetic protein circuits, containing either RELEASE or RELEASE-NOT to activate or repress protein secretion, respectively. Incubation of ASV reduced the activity of HCVP, which resulted in a reduction in the absolute amount of protein secreted in the RELEASE circuit. In comparison, when ASV was incubated with cells containing the RELEASE-NOT circuit we observed an increase in the baseline. Each dot represents a biological replicate in **g, i, and j**. Mean values were calculated from 4 biological replicates in **b** and **c**. The error bars represent +/- SEM. The results are representative of at least two independent experiments; significance was tested using an unpaired two-tailed Student’s *t*-test between the two indicated conditions for each experiment. For experiments with multiple conditions, a one-way ANOVA with a Tukey’s post-hoc comparison test was used to assess significance.\**p* < 0.05, \*\**p* < 0.01 \*\*\**p* < 0.001. \*\*\*\**p* < 0.0001.

To optimize the single-transcript design within the packaging limits of lentiviral vectors, we designed a polycistronic construct encoding a puromycin-resistance cassette, a mCherry reporter protein, a rapalog-inducible split HCVP, and secNluc-RELEASE all under the control of the EF1a promoter (**Supplementary Fig. 11a**). We validated the circuit by generating stable HEK293 cell lines via lentiviral transduction and measured secNluc, a secreted reporter protein in response to rapalog induction (**Supplementary Fig. 11b**). To optimize the dynamic range of the circuit we modified the subcellular localization of the split protease components (**Supplementary Fig. 11c**) and used the less efficient HCVP-inducible RELEASE construct^22^. These modifications increased the dynamic range, by reducing the baseline activation in the uninduced state (2.26-fold vs. 6.14-fold). Furthermore, to highlight the composable capabilities of synthetic protein circuits, we replaced the puromycin-resistance cassette with secNluc-RELEASE-NOT, enabling stable HEK293 cells to regulate multiple secreted proteins with antagonistic behaviors, via the activation of a single protease (**Fig. 4b**). In this compact design, the presence or absence of rapalog resulted in the secretion of SEAP or secNluc, respectively, without the need of additional processing components.

To assess the potential of the single-transcript design *in vivo*, we stably integrated the circuit into ARPE-19 cells via lentiviral transduction. ARPE-19 cells were selected because they are nontumorigenic, exhibit contact inhibition^68^, and have been previously engineered for cytokine delivery *in vivo*^69^. We first validated circuit performance *in vitro*, showing that engineered ARPE-19 cells activated or repressed protein secretion in response to rapalog induction via RELEASE (**Fig. 4c - left**), or RELEASE-NOT (**Fig. 4c - right**), respectively. For *in vivo* studies, we encapsulated engineered ARPE-19 cells in alginate-based microparticles and transplanted them subcutaneously into nu/nu immunocompromised mice (**Fig. 4d**). Twelve days post-transplantation, we administered rapalog (10 mg/kg) retro-orbitally and monitored secNluc secretion via *in vivo* imaging (**Fig. 4e**). Within 12 hours, we observed a significant increase in average radiance relative to animals administered a PBS control, which returned to baseline by 24 hours (**Fig. 4f**). We also observed a significant increase in circulating secNluc levels in rapalog-injected animals compared to the PBS control (**Fig. 4g**). After 25 days post transplantation, we explanted the microcapsules and observed that the encapsulated ARPE-19 cells were mCherry^+^ (**Supplementary Fig. 12a**), confirming continued circuit expression and rapalog inducibility (**Supplementary Fig. 12b**).

While we have shown the potential of using lentiviral vectors to stably engineer cells with synthetic protein circuits to control protein secretion (**Fig. 4b,c**), delivering genes using AAVs offers additional advantages such as their low immunogenicity^70^, their non-integrating nature, and tissue specificity^71^. Instead of re-optimizing the protein circuit components (**Fig. 4d**), we sought to remove non-critical gene components (i.e. puromycin and mCherry cassettes) to fit within the packaging limit of AAVs. We then incorporated a FLAG epitope tag onto the C-terminal half of split HCPV (**Supplementary Fig. 13**), which allowed us to confirm the delivery of the synthetic protein circuit using immunofluorescence. Finally, we used a smaller modified woodchuck hepatitis virus posttranscriptional regulatory element (WPRE3SL)^72^ to maximize transgene expression while reducing overall transgene size.

The optimized circuit (∼4.5 kb, **Supplementary Fig. 13**) was successfully packaged into AAVs and used to transduce primary rat cortical neurons (**Fig. 4h**), enabling rapalog-dependent control of secNluc secretion via RELEASE (**Fig. 4i**) or RELEASE-NOT (**Fig. 4j**). To improve the dynamic range, we tuned HCVP activity through co-incubation with ASV, effectively lowering the baseline secretion in RELEASE (**Fig. 4i**). However, as expected, reducing HCVP activity also decreased the dynamic range of protein secretion in RELEASE-NOT (**Fig. 4j**), a trend that remained consistent when transducing neurons with different MOIs (**Supplementary Fig. 14**). Together, these results highlight the potential of primary cell engineering using polycistronic genes with a single open reading frame that encodes synthetic protein circuits delivered via viral vectors.

### Selective expression of protein payloads using mRNA delivery

In addition to stable integration, entire synthetic protein circuits or components^73,74^ can be delivered as mRNA transcripts without any risk of insertional mutagenesis^75^. Since mRNA expression is transient^76^, RELEASE and its payload would eventually degrade, enabling repetitive and on-demand dosing. Leveraging the proteolytic requirement of RELEASE to control protein secretion, we hypothesized that we could selectively express mRNA-encoded protein payloads in engineered cells (e.g. those used in cell therapies) harboring the cognate protease, without activating RELEASE in wildtype cells (**Fig. 5a**).

**Figure 5:**
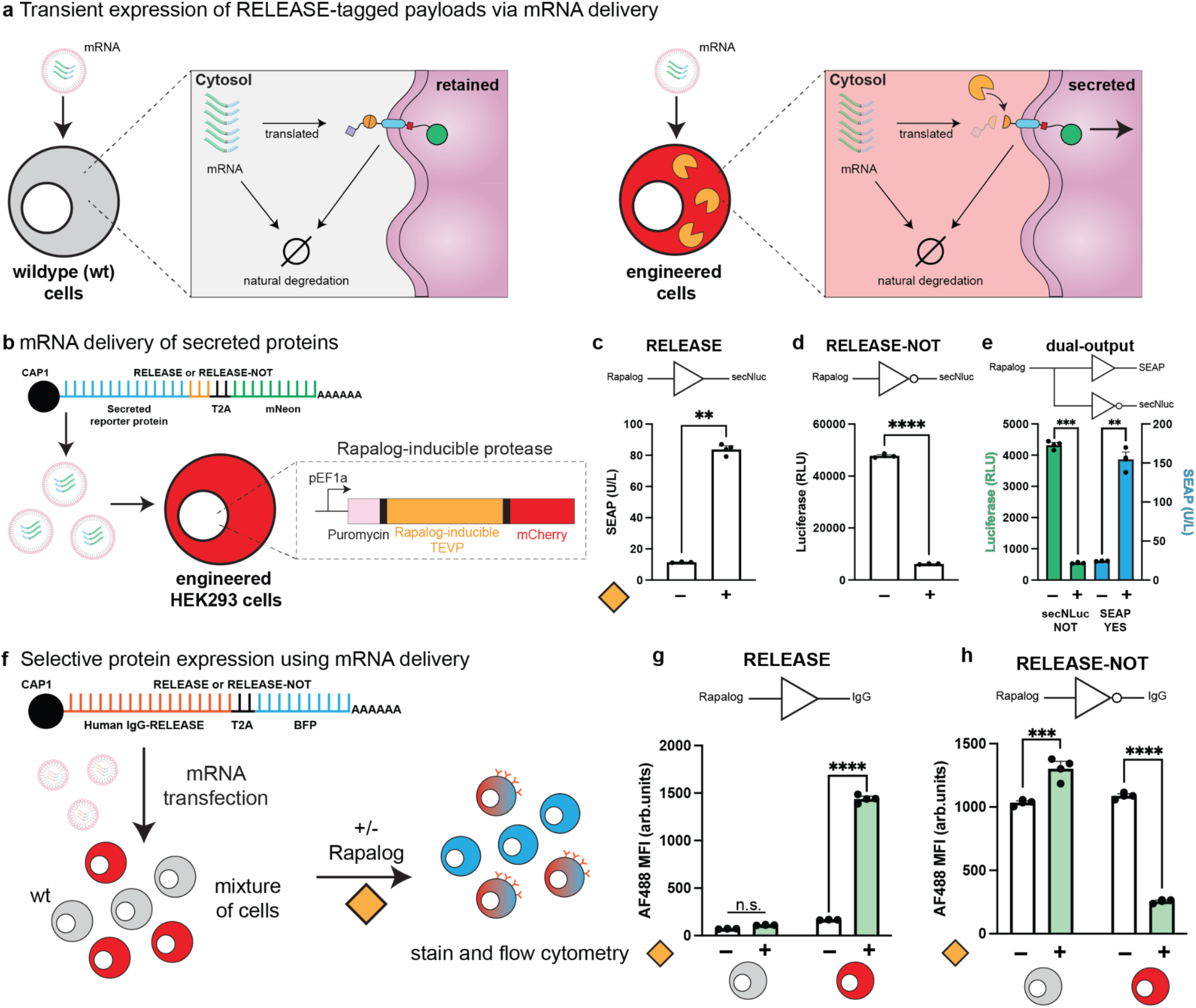
Selective expression of protein payloads using mRNA delivery. **a)** Schematic of RELEASE-tagged protein payloads delivered as mRNA. **b)** mRNA delivery of secreted proteins tagged to RELEASE or RELEASE-NOT into a rapalog-inducible TEVP HEK293 reporter cell line. **c)** RELEASE was compatible with mRNA delivery with a significant increase in SEAP secretion with rapalog induction. **d)** RELEASE-NOT was also compatible with mRNA delivery, and a significant reduction in secNLuc activity was observed with rapalog induction. **e)** Bicistronic mRNA transcript encoding both SEAP-RELEASE and secNluc-RELEASE-NOT behaved as expected with and without rapalog induction. **f)** mRNA delivery of IgG-RELEASE into a mixed population of wildtype and reporter HEK293 cells. Cells were gated based on mCherry^-^BFP^+^ and mCherry^+^BFP^+^ to measure surface expression in wildtype and reporter HEK293 cells, respectively. **g)** Wildtype cells were unable to process RELEASE, and no significant difference in IgG surface expression was observed. In comparison, engineered HEK293 cells containing the rapalog-inducible TEVP showed a significant increase in the surface expression of IgG, following rapalog induction. **h)** Surface expression of IgG using RELEASE-NOT was only observed in the reporter HEK293 cell line. Each dot represents a biological replicate. Mean values were calculated from three-four replicates. The error bars represent +/- SEM. The results are representative of at least two independent experiments. For experiments with two indicated conditions, significance was tested using an unpaired two-tailed Student’s *t*-test. For experiments with multiple conditions, significance was tested using a two-way ANOVA with a Bonferroni’s multiple comparisons test.\*\**p* < 0.01, \*\*\**p* < 0.001, \*\*\*\**p* < 0.0001.

We first tested the mRNA delivery of RELEASE-tagged secreted proteins into engineered HEK293 reporter cells containing a rapalog-inducible TEVP and mCherry (**Fig. 5b**). To do this, we first created mRNA transcripts encoding either SEAP-RELEASE or secNluc-RELEASE-NOT and the mNeon reporter separated by a T2A peptide (**Fig. 5b**). The mNeon reporter was used to visually confirm mRNA transfections before harvesting the supernatant. As expected, RELEASE (**Fig. 5c**) and RELEASE-NOT (**Fig. 5d**) showed inducible activation and repression of protein secretion, respectively. Additionally, we delivered a bicistronic mRNA transcript encoding both RELEASE and RELEASE-NOT, separated by a 2A peptide, enabling antagonistic control of two distinct secreted payloads depending on the presence or absence of rapalog (**Fig. 5e**).

Our next goal was to validate the selectivity of protein secretion in a mixed population of wildtype and engineered cells (**Fig. 5f**). We used the same bicistronic design as above, except that we encoded either IgG-RELEASE or IgG-RELEASE-NOT with the BFP reporter protein (**Fig. 5f**). The BFP reporter was selected because it was compatible with immunostaining of surface IgG. We then transfected a mixture of wild-type and engineered HEK293 cells and assessed IgG surface expression by gating cell populations based on BFP and mCherry expression using flow cytometry. From the mixture of cells, only engineered HEK293 cells (BFP^+^mCherry^+^) showed a significant increase in the surface expression of IgG when induced with rapalog (**Fig. 5g**). In comparison, wildtype HEK293 cells (BFP^+^mCherry^-^) showed comparable surface expression to uninduced engineered cells (**Fig. 5g**). Accordingly, mRNA delivery of RELEASE-NOT into a mixed population of cells only showed payload repression in the protease-high condition (**Fig. 5h**). This work highlights the use of mRNA to deliver RELEASE-tagged protein payloads, with the selectivity modulated by proteases expressed in engineered cells.

### Boolean logic integration enables conditional CAR activation

Up to this point, we have focused on expanding the RELEASE platform with single input programming capabilities and its delivery for cell or gene therapies. However, integrating multiple inputs to control protein secretion is also important for creating therapies that face more complex conditions. In our previous work^22^, we determined the design principles to implement OR- and AND-gate logic integration directly into RELEASE to control protein secretion. Now that we could implement NOT logic with RELEASE-NOT (**Fig. 1f**), our goal was to expand the programmable capabilities of RELEASE to encompass all eight possible two-input gates. By combining the design principles of the three basic gates (OR, AND, and NOT), we engineered and validated the remaining Boolean logic gates using TEVP and HCVP as inputs (**Fig. 6a**). For certain gates, such as XOR and NAND, achieving the expected behavior required co-transfection of two different RELEASE variants. All designs successfully exhibited their intended logic functions on protein secretion (**Fig. 6a**).

**Figure 6:**
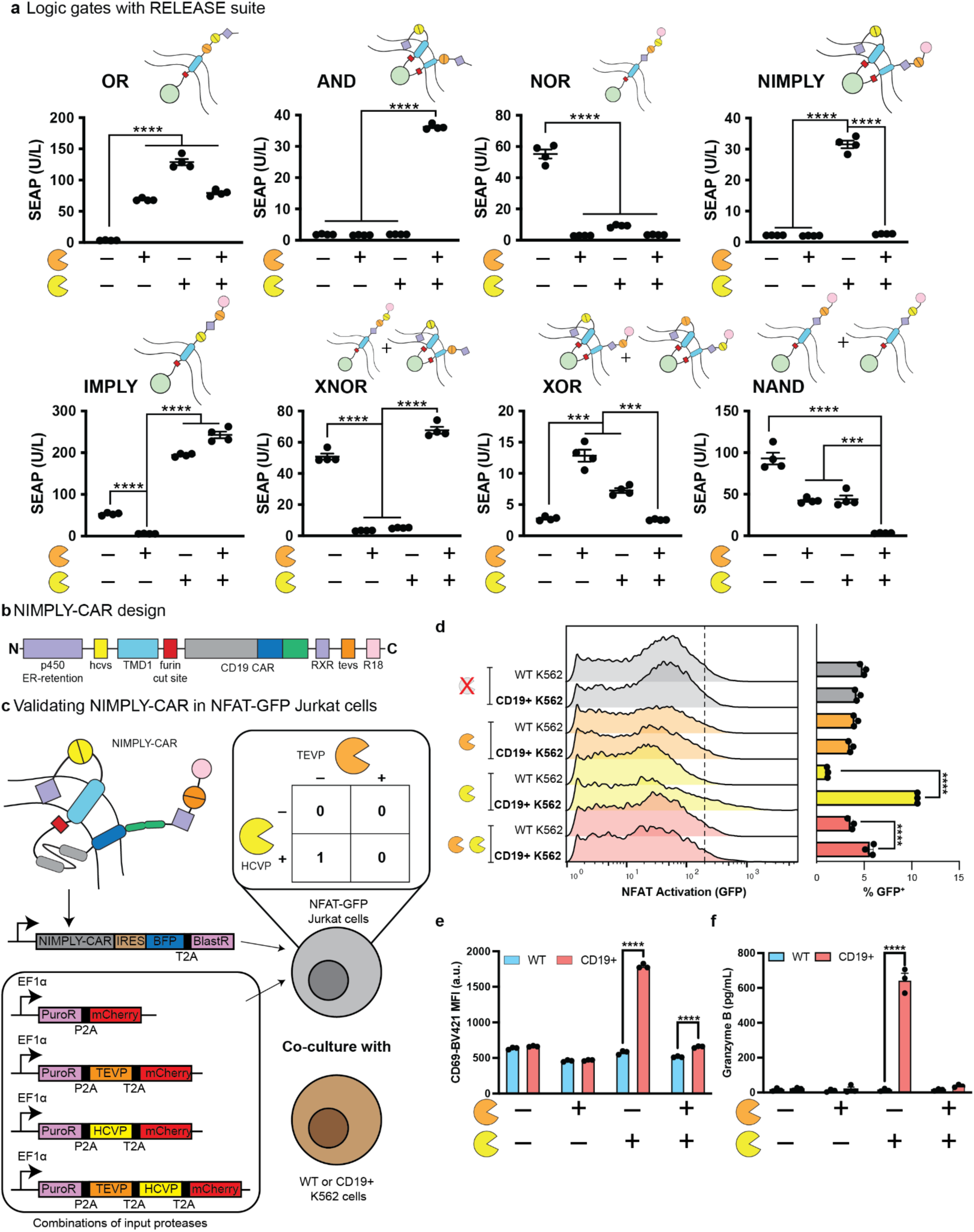
Boolean logic integration enables conditional CAR activation. **a)** RELEASE variants were engineered to enable functional completeness of Boolean logic gates. For each gate, there is a schematic representation of the respective RELEASE variant to confer its behaviour. TEVP and HCVP served as binary inputs, which were either included or excluded in the transient transfections. SEAP secretion served as an output. **b)** Schematic representation of NIMPLY-CAR design used to conditionally express anti-CD19 CAR using RELEASE-NIMPLY in response to HCVP expression (yellow pac-man). **c)** Schematic representation of the co-culture assay designed to validate the NIMPLY-CAR behavior. NIMPLY-CAR was stably integrated into NFAT-GFP Jurkat cells via lentiviral transduction. From this original cell line, four additional NFAT-GFP cell lines were generated, each containing an additional transgene encoding one input condition of all possible two-input protease combinations. Engineered NFAT-GFP Jurkat cells were co-cultured with either wildtype or CD19^+^ K562 cells. **d)** Representative histograms of GFP expression (NFAT activation) for each possible combination of co-culture conditions. The dotted line represents the threshold for GFP expression relative to a untransduced NFAT-GFP Jurkat cell control. There was a significant difference in the percentage of GFP^+^ cells in the HCVP only and the dual protease (HCVP^+^/TEVP^+^) conditions for NFAT-GFP Jurkats co-cultured with CD19^+^ K562 cells, relative to wildtype K562 cells. **e)** Average MFI for CD69+ expression was significantly increased in the HCVP and dual protease conditions in NFAT-GFP Jurkats co-cultured with CD19^+^ K562 cells, relative to wildtype K562 cells. **f)** There was a significant increase in Granzyme B secretion in the HCVP alone condition in NFAT-GFP Jurkats co-cultured with CD19^+^ K562 cells. Each dot represents a biological replicate. Mean values were calculated from three-four replicates. The error bars represent +/- SEM. The results are representative of at least two independent experiments. For the Boolean logic gates, statistical significance was calculated using a one-way ANOVA with a Tukey’s post-hoc comparison test. For every other measurement, statistical significance was calculated using a two-way ANOVA with Bonferroni’s multiple comparisons test. \*\*\**p* < 0.001. \*\*\*\**p* < 0.0001.

Since RELEASE functions independently of the payload, our next goal was to transition towards controlling functionally relevant outputs. We hypothesized that chimeric antigen receptors (CARs) would benefit from multi-input control of its surface expression, given the significant off-target effects associated with CAR-T therapy^77,78^ and their compatibility with mRNA delivery^79^. We modified the RELEASE-NIMPLY variant for the conditional surface expression of anti-CD-19 CAR (NIMPLY-CAR, **Fig. 6b**) when only HCVP was expressed. To assess logic-dependent expression we stably integrated NIMPLY-CAR and all possible two-input combinations of HCVP and TEVP (**Fig. 6c**) into a NFAT-GFP Jurkat reporter cell line^80^. We then measured NIMPLY-CAR surface expression by immunostaining for the extracellular myc-tag on the CAR and observed the greatest surface expression in the HCVP only condition; however, the TEVP only condition also showed an increase relative to the no protease control, but at a lower level than the HCVP only condition (**Supplementary Fig. 15a**).

Following expression validation of the NIMPLY-CAR design, we next tested T-cell activation in the engineered NFAT-GFP Jurkat cells by co-culturing them with either wildtype or CD19^+^ K562 cells. After 24 hours of co-culture, we observed a significant increase in GFP expression and the percentage of GFP^+^ cells in the HCVP-only condition (**Fig. 6d**), indicating T-cell activation in response to conditional CAR surface expression. Additionally, we observed a significant increase in T-cell activation in the dual protease condition, albeit a smaller increase than the HCVP-only condition, suggesting incomplete repression of surface expression (**Fig. 6d**). We observed similar trends when staining for CD69^+^ expression (**Fig. 6e**), and early T-cell activation marker^81^. Interestingly, only the HCVP-only condition resulted in significant secretion of granzyme B (**Fig. 6f**), suggesting that despite some leakiness in the NIMPLY-CAR design (**Fig. 6d,e**), the level of surface expression in the dual protease condition was insufficient to trigger full cytotoxic effector function. We also observed a significant increase in IL-2 secretion in the HCVP-only condition when co-incubated with CD19^+^ K562 (**Supplementary Fig. 15b**). These results highlight the ability to encode Boolean logic directly into the regulation of both secreted and surface-expressed therapeutically-relevant proteins using RELEASE, streamlining the circuit architecture by minimizing the need for intermediate proteases.

## Discussion

Here, we introduce the RELEASE suite, a collection of engineered proteins with expanded input-processing capabilities, enabling complete control over intercellular signals in mammalian cells. This complete suite supports a range of advanced functionalities, including negative repression of protein secretion (**Fig. 1f**), tunable input sensitivity (**Fig. 2b,c**), control over payload magnitude (**Fig. 3**), quantitative processing (**Fig. 2g, i**), and multi-input logic (**Fig. 6a**), all without requiring intermediate proteases. Because the RELEASE suite is highly modular and functions independently of the payload, we demonstrated conditional activation of CAR, without the need for re-optimization (**Fig. 6b-f**). Moreover, the simplified architecture of the RELEASE suite enables compatibility with clinically relevant delivery modalities, including viral vectors (**Fig. 4**), and mRNA delivery (**Fig. 5**), both *in vitro* and *in vivo*. Together, this complete suite lays the foundation for programming sophisticated cellular behaviors with therapeutic potential, entirely at the protein level.

With an expanded RELEASE suite, numerous additional programmability has been achieved with minimal increases of the genetic footprint. This is in large part due to the reliance on ubiquitously expressed native proteins, such as endoproteases and 14-3-3 scaffolding proteins, which are expressed in all eukaryotic cells. By leveraging these endogenous mechanisms for subcellular protein translocation, we anticipate these tools being compatible across any eukaryotic cell lineage, including those from different species. Nevertheless, it is important to acknowledge that differential expression of 14-3-3 proteins^82^ in specific cells and tissues may influence the absolute levels of protein secretion or surface expression when using RELEASE variants that rely on 14-3-3 proteins for their operations (**Fig. 1f**, **Fig. 2c, g, i, and Fig. 6a**). There is some level of tuning that can be done by using different TMDs with RELEASE-NOT (**Supplementary Fig. 10**), but fundamentally the maximum secretion capacity is directly dependent on the reversible binding of 14-3-3 to the R18 peptide present at the c-terminus. Optionally, overexpression of the 14-3-3ζ isoform may normalize these potential differences (**Supplementary Fig. 16**); however, this would increase the construct size and has the potential for off-target binding. Although we’ve only focused efforts on utilizing these two endogenous mechanisms in RELEASE, it is conceivable that other endogenous mechanisms or even additional post-translational modifications may be leveraged to further optimize and expand the potential of synthetic protein circuits using RELEASE.

By utilizing high-throughput screening techniques, we were able to characterize a wide range of protein expression (**Fig. 3c-e**) and identify two broad classes of transmembrane domains for use in RELEASE: inducers and non-inducers (**Fig. 3c**). We hypothesize there is the opportunity for more optimization of these TMDs by altering the juxtamembrane or hinge residues included. Furthermore, we see utility in expanding this screening platform to optimize multi-pass RELEASE architectures to find optimal combinations of TMDs and for use in assessing the specific dynamics of RELEASE activation. Screening combinatorial TMD pairings could improve understanding of multi-membrane receptors^83^ and expand optimization efforts of applications such as NIMPLY-CAR, which utilize a dual-membrane architecture (**Fig. 6c**). Nevertheless, we see the current screen as a powerful resource for the fields of synthetic receptor engineering or controlled secretion, as surface display and secretion correlations were well preserved regardless of gene delivery method, protein output, and cell type (**Fig. 3i-l**).

One of the longstanding challenges in translating synthetic protein circuits into therapeutic contexts has been efficient and stable delivery. To date, most *in vivo* demonstrations of protein circuits have relied on DNA transient transfections^24,25,28^, limiting their relevance for translatable applications. To address this, we leveraged the compact nature of post-translational protein circuits to develop single-promoter polycistronic designs compatible with clinically relevant delivery vectors, such as lentivirus and AAVs (**Fig. 4d, h**). Unlike multi-promoter designs, which can suffer from stochastic silencing of individual promoters resulting in unpredictable behavior^84,85^, a single-promoter construct ensures that the entire circuit is either active or silenced, which may improve safety in therapeutic settings. In addition, synthetic protein circuits can synergize with existing transcriptional systems to provide an orthogonal level of control. This is particularly advantageous when using synthetic promoters or enhancers tuned to specific cell types^86,87^ or states^88^ that may have inherently weak activity, limiting their utility for some biomedical applications. The use of RELEASE-based circuits could compensate for this by tuning circuit outputs through altering the TMDs (**Fig. 3**) or protease cut site kinetics (**Fig. 2**). However, it should be noted that this approach would only be amenable for tuning secreted or surface expressed outputs, and the choice of TMD may require additional optimization depending on the application.

Using RELEASE to fine tune secreted or surface expressed outputs offers additional advantages as well. Protein circuits are compatible with mRNA delivery, which do not use promoters to control therapeutic outputs. By swapping TMDs and leveraging their effect on post-translational control (**Fig. 3c**), we can precisely modulate levels in ways previously not possible for secreted outputs. This, combined with the selectivity offered by delivering RELEASE via mRNA (**Fig. 5f**) would be particularly advantageous for therapeutic applications where tight control over cytokine or biologic dosing is critical to avoid systemic toxicity or off-target effects. Notably, mRNA delivery of CARs has already been explored clinically as a transient and safer alternative to viral-engineered T cell therapies^79^, demonstrating the feasibility of applying mRNA to deliver our NIMPLY-CAR platform (**Fig. 6d**) for conditional control. Although we did not deliver RELEASE via lipid nanoparticles (LNPs), others have successfully used LNPs to deliver synthetic protein circuits for cancer therapy^74^. Given the modular, plug-and-play nature of synthetic protein circuits, we anticipate seamless integration with LNP-based delivery in future studies.

Another limitation of our current platform is the reliance on non-human proteases to activate the RELEASE suite, which could potentially be immunogenic in therapeutic applications. In future work, one could address this by implementing human-derived proteases or engineering human proteases with altered substrate specificities to create orthogonal systems^73^. Importantly, because the RELEASE suite enables the programming of complex behaviors, such as single-input, multi-output secretion (**Fig. 4b**) and quantitative input processing (**Fig. 2g, i**) using a single protease input, the burden of engineering multiple orthogonal proteases is considerably reduced. Overall, by enabling control across multiple layers of the central dogma, the RELEASE suite paves the way for the design of more efficient, compact, and programmable cellular and gene therapies capable of executing increasingly sophisticated behaviors in complex biological environments.

## Materials and Methods

### Plasmid generation

All plasmids were constructed using general practices. Backbones were linearized via restriction digestion, and inserts were generated using PCR, or purchased from Twist Biosciences. WPRE3-SVLpA sequence for AAV vector generation purchased from ANSA Biotechnologies as the sequence was not able to be synthesized elsewhere. All new plasmids used in this study will be deposited with annotations to Addgene (https://www.addgene.org/Xiaojing_Gao/).

### Tissue culture

Human Embryonic Kidney (HEK) 293 cells (ATCC, CRL-1573) and HEK293T-LentiX (Takara Biosciences) were cultured in standard culture conditions (37°C and 5% CO2) in Dulbecco’s Modified Eagle Media (DMEM), supplemented with 10% Fetal Bovine Serum (FBS, ThermoFisher; catalog# FB12999102), 1X Pen/Strep (Genesee; catalog# 25-512), 1X non-essential amino acids (Genesee; catalog# 25-536), and 1X sodium pyruvate (Santa Cruz Biotechnology; catalog# sc-286966). ARPE-19 cells (ATCC, CRL-2302) were cultured in standard culture conditions in DMEM/F-12 (1:1) GlutaMAX media (ThermoFisher; catalog# 10565018), supplemented with 10% FBS, and 1X PenStrep. K562 cells (ATCC, CCL-243) were cultured in RPMI-1640 (Sigma-Aldrich; catalog# R8758) media supplemented with 10% FBS, and 1X PenStrep. Primary Rat Cortical Neurons (Gibco, catalog# A36512) were cultured in Complete Neurobasal™ Plus Medium (Gibco; catalog# A3582901) in a 24 well-plate coated with Poly-D-Lysine (4.5 µg/cm^2^). Both wildtype and NFAT-GFP Jurkat cells were cultured in RPMI-1640 (Sigma-Aldrich; catalog# R8758) media supplemented with 10% heat-inactivated FBS, and 1X PenStrep. HEK293 cells were used for all transient transfection experiments and K562 cells were used for the high-throughput transmembrane domain library. HEK293T cells were used to produce lentivirus, described below. All cells tested negative for mycoplasma.

### Transient transfection

HEK293 cells were cultured in 96-well tissue-culture treated plates under standard culture conditions. At 70-90% confluency, cells were transiently transfected with plasmids using the jetOPTIMUS® DNA transfection Reagent (Polyplus transfection; catalog# 117-15), as per manufacturer’s instructions. A complete list of plasmids used in each figure with their respective amounts can be found in **Supplementary Table 3**.

### Measuring protein secretion

Protein secretion was measured by a Secreted Embryonic Alkaline Phosphatase (SEAP) assay, as previously described^15^. Briefly, two days after transient transfection or induction with A/C heterodimerizer (Takara Biosciences; catalog# 635056, referred to as rapalog for the rest of the manuscript), the supernatant was collected and heat-inactivated at 70°C for 45 minutes. After heat-inactivation, 10 - 40 mL of supernatant was mixed with dH_2_O to a final volume of 80 mL. This mixture was then mixed with 100 μL of 2X SEAP Buffer (20 mM homoarginine (ThermoFisher catalog# H27387), 1 mM MgCl2, and 21% (v/v) diethanolamine (ThermoFisher, catalog# A13389)) and 20 μL of the p-nitrophenyl phosphate (PNPP, Acros Organics catalog# MFCD00066288) substrate (120 mM). Samples were measured via kinetic measurements (1 reading/minute) for a total of 30 minutes at 405 nm using a SpectraMax iD3 spectrophotometer (Molecular Devices) with the Softmax pro software (version 7.0.2)

Secreted NanoLuc® (secNLuc) was measured by taking the supernatant from samples that were previously transiently transfected or induced with rapalog. Briefly, 5 µL of supernatant was mixed with 45 µL of dH2O and then mixed with 50 µL of the Nano-Glo® Luciferase Assay system (Promega; catalog# N1110), per the manufacturer’s instructions. Samples were measured using a SpectraMax iD3 spectrophotometer (Molecular Devices) with the Softmax pro software (version 7.0.2).

### Lentiviral transduction for cell engineering

To generate lentivirus, HEK293T-LentiX cells (Takara Biosciences) were transfected with 750 ng of an equimolar mixture of the three third-generation packaging plasmids (pMD2.G, pRSV-Rev, pMDLg/pRRE) and 750 ng of donor plasmids using 200 uL of jetOPTIMUS® DNA transfection Reagent (Polyplus transfection, catalog# 117-15). pMD2.G (Addgene plasmid #12259, https://www.addgene.org/12259/), pRSV-Rev (Addgene plasmid #12253, https://www.addgene.org/12253/), and pMDLg/pRRE (Addgene plasmid # 12251, https://www.addgene.org/12251/) were gifts from Didier Trono. For some experiments, we generated lentivirus using the second-generation packaging plasmids (600 ng of PAX2, 300 ng of pMD2g and 1,100 ng of the transfer plasmid). After 1 - 3 days of incubation, the lentivirus was harvested and filtered through a 0.45 mm filter (Millipore) to remove any cellular debris. The harvested virus was precipitated using the Lentivirus Precipitation Solution (Alstem; catalog# VC100) and centrifuged at 1,500 x g at 4°C for 30 minutes to remove any residual media. The concentrated virus was then resuspended in supplemented DMEM or RPMI, aliquoted and frozen at −80C for later use.

Lentivirus was titrated on HEK293, ARPE-19, or Jurkat cells by serial dilution in supplemented DMEM, DMEM/F-12, or RPMI-1640 media respectively. Depending on the cell line, 72 hours after incubation the percentage of mCherry^+^ or BFP^+^ cells were quantified using flow cytometry. For some cell lines, the cells were selected using 1000 ng/mL of puromycin (ThermoFisher Scientific; catalog# J61278-MB) for one week until >90% of cells were mCherry-positive before being used for *in vitro* or *in vivo* experiments. For mRNA transfection experiments, the transduced HEK293 reporter cells were sorted using the Stanford Shared FACS facility (RRID: SCR_017788) to generate cell lines with different expression levels of inducible TEVP.

### Adeno-associated virus (AAV) generation and transduction

The AAVs were manufactured through the Stanford Gene Vector and Virus Core (GVVC). Plasmids were grown up in 50 mL cultures of LB and then extracted using QIagen plasmid midiprep kit, as per manufacturer’s instructions. GVVC produced a set of AAVs with the AAV8 (with Y733F mutation) serotypes. The two AAVs were measured for their ITRs: AAV8-CC593: 1.80 ×10^13^ vg/mL, and AAV8-CC594: 1.97 ×10^13^ vg/mL. The two AAV8 constructs were used to transduce rat cortical neurons at different multiplicities of infection (MOI) ranging from 12,500 to 125,000 in 1 mL of Neurobasal Media (Gibco, catalog# 21103049) per well. After 7 days of culture, 300 µL of media was replaced with media containing either Rapalog or vehicle to a final concentration of 200 nM. Cells were incubated for 2 days before harvesting supernatant to measure secNluc secretion. For samples incubated with Asunaprevir (ASV, Fisher Scientific, catalog# NC1026214) at a final concentration of 100 nM.

### Rat Cortical Neuron Staining

Following secretion experiments, transduction was validated by antibody staining. Media was aspirated, and rat cortical neurons were were washed three times with 500 µL with DPBS without Ca or Mg (GenClone, Cat #: 25-508). DPBS was allowed to sit on the cells for 3 minutes between washings. were fixed using 500 µL paraformaldehyde (PFA) (Thermo 16% PFA catalog 28908, 4% in DPBS) for 30 minutes. PFA was removed from the cells and residual PFA was removed with another 500 µL wash of DPBS. Neurons were then permeabilized for 5 minutes using 0.3% Triton X in DPBS. Neurons were then washed three times with PBS as before. APC anti-DDDDK primary antibody (abcam, catalog# ab72569 Lot: 1029635-7) was diluted 1:500 in DBPS + 5% bovine serum albumin (BSA) and 500 µL was placed on the neurons and allowed to incubate overnight at 4°C without light. Antibody stain was aspirated off of the cells and cells were washed three times with 500 µL DBPS to remove residual antibody. Stained neurons were then coated with 300 µL of DAPI-containing mounting buffer (Cell Signaling Tech. Catalog #8961S) and allowed to incubate in the dark for 5 minutes before collecting images using EVOS M7000 using DAPI (AMEP4950) and Cy5.5 (AMEP4973) light cubes. 5 images were collected per well using EVOS automatic image collection program.

### Alginate microencapsulation

ARPE-19 cells were trypsinized and centrifuged at 300 x g for 7 minutes, and then resuspended in a minimal volume of PBS. The cell/PBS slurry was mixed with a sodium alginate solution (Novamatrix, 4%wt in PBS) using a female-female mixing elbow and two 1-mL luer lock syringes to reach the desired alginate concentration of 1.7%wt and 2.0 ×10^6^ cells/mL. A concentric needle (inner diameter = 0.584 mm, outer diameter = 1.10 mm) was affixed to the end of the syringe and placed onto a syringe pump. Compressed air was controlled using a calibrated flowmeter and flowed through the outer nozzle at a flow rate of 1.5 standard litres per minute. The alginate solution was extruded into a bath of 50 mM SrCl_2_ (Sigma Aldrich, catalog# 255521) and 200 mM mannitol (Sigma Aldrich, catalog# M4125) at a flow rate of 125 mL/min. The particles were collected using a 70 µM filter (Celltreat) and rinsed three times with PBS. Encapsulated cells were incubated overnight in standard culture conditions (37°C and 5% CO2) in supplemented DMEM/F12. The microcapsules were then back loaded into a 1 mL syringe, containing 400 μL of 0.75%wt hydroxypropyl methylcellulose (HPMC – Sigma Aldrich, catalog# H3785) in PBS and gently mixed by moving the barrel of the syringe up and down until the solution was homogenous.

### Cell transplantation and chemical induction of protein secretion

All animal studies were performed in accordance with the National Institutes of Health (NIH) guidelines, with the approval of the Stanford Administrative Panel on Laboratory Animal Care. Encapsulated ARPE-19 cells were subcutaneously injected into the right flank of nu/nu mice (Charles River, Crl: NU-*Foxn1^nu^*). For induction of the engineering cells, rapalog (5 mg, Takara Biosciences) was first reconstituted in *N,N*-Dimethylacetamide (Sigma-Aldrich) to make a 10 mg/mL stock solution, as previously described^89^. Prior to induction, the rapalog solution was mixed with equal parts of a PEG400:Tween 80 (9:1 ratio), and then animals were randomly assigned into different groups and administered either the rapalog solution, or vehicle alone via intraperitoneal injection (i.p.).

### In vivo imaging and analysis

To measure secNLuc, mice were injected i.p. with 0.546 mg fluorofurimazine (Selleckchem, catalog:S5565) solubilized in sterile filtered water with Pluronic F-127 as previously described^90^. After 10 minutes, mice were anesthetized with isoflurane and a broadband luminescent reading was taken with an exposure time of 30s (PerkinElmer IVIS Lumina III). The signal was quantified as the average radiance in the region of interest, which was defined as a circle of consistent size around the subcutaneous injection site (Living Image software, PerkinElmer). Luciferase measurements were taken before administering rapalog and 4, 12, and 24 hours post-injection with rapalog.

### Human transmembrane domain library design

Annotated human transmembrane domains were queried from the Uniprot Database^91^, and each element of the library was a single spanning transmembrane domain. To select type I transmembrane domains, we retrieved all transmembrane domains from proteins that contained a signal peptide sequence. For multipass transmembrane proteins, each type I transmembrane domain was treated as an individual element, while the type II transmembrane domains from these proteins had their sequences reversed to adopt a pseudo-type I orientation and were included in the library. For each type I transmembrane domain, we also retrieved the native juxtamembrane domains (+8 amino acids from the c-terminal of the transmembrane domain). For library elements containing type II transmembrane domains, since their natural juxtamembrane domains would be extracellular-facing, we added the juxtamembrane domain of CD8a to each element (i.e. -LYCNHRNR). In addition, we selected a random small subset of transmembrane domains from multi-spanning membrane proteins that do not insert via a signal peptide mechanism. Finally, we also included 5 human transmembrane domains commonly associated with immunoengineering: CD4 (Uniprot ID: P01732), CD8 (Uniprot ID: P01730), CD3 (Uniprot ID: P04234), and CD25 (Uniprot ID: P01589). For library elements associated with these specific domains, we performed an alanine scan, sequentially included different numbers of amino acids in the juxtamembrane, tiled the transmembrane domain in 37 amino acid windows, and circular permutations of the transmembrane domains (**Supplementary Table 2**). Duplicate sequences were removed, amino acid sequences were reverse-translated, and codon optimization was performed for human codon usage (reverse translation and DNA optimization was performed using DNA chisel^92^). 400 random controls of 63 base pairs without stop codons were computationally generated to act as negative controls.

### Human transmembrane domain library cloning

A Twist oligo pool was reconstituted in 10 µL of nuclease-free water and the library was PCR amplified using primers specific to adapters flanking the sequence. To reduce contamination, all reactions were performed in a pre-PCR hood. A test PCR was performed using 1 µL Hercules II Fusion DNA polymerase (Agilent, catalog# 600675), with 0.1 µL of the oligo pool, 0.2 µL of each 10 µM amplification primers, 1 µL DMSO, 10 µL of Herculase 5X buffer, and nuclease-free H_2_O to 50 µL. The PCR protocol was 3 minutes at 98°C, followed by cycles of 98°C for 20 seconds, 58°C for 20 seconds, 72C for 30 seconds, and a final extension step at 72°C for 3 minutes. The number of cycles used to amplify the library was determined as the minimum number of cycles required to see a clean visible DNA product for extraction, which was 10 cycles for this library. The amplified dsDNA (expected size of ∼300 bp) was loaded into 6 wells of a 2% TAE gel and purified using a QIAgen gel extraction kit (QIAgen). The library was cloned into a lentiviral RELEASE backbone (EF1a_hIgG1-LCS-tevs-RXR6.2-T2A-BFP-P2A-Blast) using 24 x 10 uL GoldenGate reactions (75 ng of pre-digested and purified backbone, 5 ng of amplified library, 2 µL of 10X T4 DNA ligase buffer (NEB), and NEB GoldenGate BsmBI-V2 assembly kit) with 30 cycles of digestion at 37°C and ligation at 16°C for 5 minutes each, followed by a final 5 minute digestion at 37°C and then 20 minutes of heat inactivation at 70°C. The reactions were pooled and then eluted using two DNA clean and concentrator columns (Zymo Research, catalog# D4033) so that both elutions had a final volume of 6 µL. 2 µL of the pooled GoldenGate reactions per tube were transformed with 50 µL of Endura electrocompetent cells (Lucigen, catalog# 60242-2) as per manufacturer’s instructions (BioRad Gene Pulser Xcell, 1.8 kV, 600 Ohm, 10 µF). After recovery, the transformed cells were plated on 4 large 10” x 10” plates containing LB agar supplemented with carbenicillin and incubated overnight at 37°C. In parallel, 4 small plates were prepared using diluted transformed cells to count colonies and confirm that we had at least >50X library coverage. After overnight incubation, 2 of the larger plates were scraped together and the plasmid pools were extracted using a HiSpeed Plasmid Maxiprep kit (QIAgen). To characterize the quality of our library, the transmembrane domains were amplified using primers with extensions that included the Illumina adapters, which were then sequenced.

### Inducible expression assay and magnetic separation

To generate the reporter cell line, K562 were electroporated in Amaxa solution (Lonza Nucleofector 2b, setting T0-16) with 1µg of reporter donor (pEF1a-p450-rapalog-split-TEVP-T2A-mCherry) and 1 µg of SuperPiggyBac transposase. After 7 days, the cells were treated with 1000 ng/mL of puromycin for two weeks to ensure that the construct was stably integrated. A monoclonal population was generated through serial dilution. The expression of the fluorescent reporter was measured by microscopy and flow cytometry. The SuperPiggyBac transposase plasmid was a gift from the Bassik lab. A large-scale lentiviral production and spinfection was performed to transduce the human transmembrane domain library into the reporter K562 cell line, as previously described^60^. Briefly, 9 x 15-cm dishes were each plated with 1 ×10^6^ HEK293T-LentiX cells in 10 mL of supplemented DMEM media, grown overnight, and then transfected with an equimolar mixture of the three third-generation packaging plasmids (pMD2.G, pRSV-Rev, pMDLg/pRRE) and 4 µg of donor plasmids using 1 mL of jetOPTIMUS® DNA transfection Reagent (Polyplus transfection, catalog# 117-15). pMD2.G (Addgene plasmid #12259, https://www.addgene.org/12259/), pRSV-Rev (Addgene plasmid #12253, https://www.addgene.org/12253/), and pMDLg/pRRE (Addgene plasmid # 12251, https://www.addgene.org/12251/) were gifts from Didier Trono. After 48 hours the virus was harvested, pooled, and filtered through a 0.45-µM PVDF filter (Milipore) to remove any cellular debris. For the human transmembrane domain screen, 2.25 x 10^7^ K562 reporter cells were transfected via spinfection for 2 hours, with two biological replicates for the infections. Following 3 days after infection, infected cells were cultured with 10 µg/mL of blasticidin (Sigma Aldrich). The infection and selection efficiency was periodically checked using flow cytometry by measuring BFP. Cells were maintained in log growth conditions in a T-225 flask by diluting cell concentrations back to 5 x 10^5^ cells/mL, with at least 1.5 x 10^7^ cells per replicate to ensure sufficient library coverage (at least 2500 cells per element) and minimize potential element dropouts. On day 12 post-infection, each bio-replicate was separated into two groups (∼2.5 x 10^7^ cells each), where one of the groups was administered 300 nM of rapalog and the other an equal volume of the vehicle for 3 days. Following 3 days of incubation, we had a library coverage of >10,000 per element and cells were harvested for magnetic separation.

Magnetic separation was performed as previously described^60^. Briefly, a synthetic cell surface marker consisting of an Igk signal sequence linked to a human IgG1 FC region conjugated to RELEASE containing the variable transmembrane domain (**Supplementary** Figure 7a). Cells were incubated with rapalog (300 nM) for 3 days to induce the surface expression of the synthetic surface marker to enable magnetic separation. Blocking buffer was prepared by supplementing PBS with 2 mM EDTA and 2% BSA, which was then filtered through a 0.22 µm filter (Millipore). Cells were spun at 300 x g for 7 minutes and the media was aspirated. The cells were washed twice using PBS to remove any residual IgG from the media. Dynabeads M-280 Protein G (Thermofisher, catalog# 10003D) were resuspended by vortexing for 30s. For each 1 ×10^7^ of cells, 90 µL of beads were prepared by mixing with 500 µL of the blocking buffer, vortexing, placing on a magnetic tube rack for 2 minutes and then reconstituted in 600 uL of blocking buffer. For each condition in the TM library, 7.5 x 10^7^ cells were magnetically separated using 675 µL (resuspended in 4.5 mL of blocking buffer), and put into a 15 mL falcon tube on a large magnetic rack. After incubation for 2 minutes, the unbound fraction was harvested and put into a new tube for a subsequent incubation on the magnetic rack to remove any residual beads. The magnetic beads were incubated with the same volume of blocking buffer for 2 minutes on the large magnetic rack, and the supernatant was discarded. The beads (representing the bound fraction) were resuspended in 1 mL of blocking buffer. Finally, the unbound and bound fractions were spun down at 300 x g for 7 minutes, and the cell pellets were stored at −20°C until genomic DNA extraction.

### Library Preparation and next-generation sequencing

Genomic DNA was harvested using a QIagen Blood Maxi Kit as per manufacturer’s instructions. For the bound fractions, following cell lysis the samples were placed on a large magnetic rack to remove the magnetic Dynabeads and ensure they did not clog the column. The harvested genomic DNA was eluted in EB and then amplified by PCR using primers containing Illumina adapters as extensions. A test PCR was performed to determine the minimum number of cycles required to see a clear band at the expected size (∼200 bp). To perform the test PCR, 5 µg of genomic DNA was mixed in a 50 µL reaction of NEB High-Fidelity 2X PCR Mastermix. Following the test PCR, 15 x 50 µL reactions were set up per fraction (5 µg genomic DNA, 0.5 µL of each 100 µM primer) with the following parameters: 98°C for 3 minutes, 21 cycles of 98°C for 10 seconds, 63°C for 30 seconds, and 72°C for 30 seconds, followed by a final extension step at 72°C for 2 minutes. The PCR reactions for each fraction were pooled and 200 µL from each fraction was run on a 2% TAE agarose gel, the library band around 300 bp was excised, and the DNA was purified using the QIagen gel extraction kit. These fractions were then quantified using a Qubit HS kit (thermofisher), pooled with 15% PhiX control (Illumina) and sequenced through Admera Health using a HiSeq 2×150 (Illumina).

### Human transmembrane domain analysis

Sequencing reads were demultiplexed using bcl2fastq^60^. Analysis of NGS sequences was conducted using a custom Python script performing an exact string match of the unique TM domain, defined as the sequence between the forward and reverse handles of the regions used to amplify sequences from the genomic sequences. Counts of each library element were calculated and normalized to the sequencing depth of the corresponding pool. Read count values were further normalized by the amount of genetic DNA collected from each subpopulation, representative of the total amount of each library element in each fraction to account for both variations in representation within each bin as well as differential separation due to magnetic separation. The enrichments for each domain were calculated between Bound and Unbound samples within each respective fractions. For elements that had less than 5 reads in both replicates, the element was assigned a value of 0 counts, while elements that had less than 5 reads within a single replicate had their reads adjusted to 5 to avoid overinflation of enrichments from low depth. The surface expression score was calculated as the log_2_(Bound:Unbound) of the ON population, and the ER retention score was calculated as the log_2_(Bound:Unbound) of the OFF population. The TEVP inducibility score was calculated as the ratio of these two enrichment scores log_2_(ON:OFF). For surface expression, or TEVP inducibility, hits were elements that had a log_2_ score that was ≥ 3 standard deviations from the mean of the surface expression or TEVP inducibility score for the 400 randomized controls, respectively.

### mRNA synthesis via in vitro transcription

Genes encoding secreted proteins under the control of RELEASE were cloned into a plasmid template flanked by optimal 5’ and 3’ UTRs containing a T7 promoter, which was a gift from Prof. Michael Elowitz. For the IgG-RELEASE constructs, the 5’ and 3’ UTRs were taken from the human alpha-globulin RNA with an optimized Kozak sequence, and the amino-terminal enhancer of split mRNA and mitochondrial encoded 12s ribosomal RNA, respectively^93^. A Poly-A-tail was added to the DNA template via PCR using the Q5 PCR master mix (New England Biosciences, M0494L) and a reverse primer containing a 200 bp polyA sequence. Following amplification, the PCR products were cleaned and concentrated using the DNA Clean & Concentrator kit (Zymo Research, catalog# D4033). In vitro transcript mRNA was synthesized from the DNA template using HiScribe IVT kit (New England Biosciences, catalog# E2040S) supplemented with 100% N1-methyl-psuedouridine-5’-triphosphate (Trilink, catalog# N-1081-10) modified bases and a murine RNase inhibitor (New England Biosciences, catalog#M0314). The synthesized mRNA was co-transcriptionally capped with a Cap 1 structure using CleanCapAG (Trilink, catalog# N-7113) and purified using Monarch RNA Cleanup kit (New England Biosciences, catalog# T2040). The synthesized mRNA was stored in −80°C until used for mRNA transfections.

### mRNA transfection

For mRNA transfections, 1 x 10^5^ HEK293 cells were plated per well on a 24-well plate. The following day, 250 ng of mRNA was transfected per well using the TransIT®-mRNA Transfection Kit (Mirus Bio, catalog# MIR2250). To induce the reporter cell lines, rapalog was administered at the time of transfection. All experiments were induced with 100 nM of rapalog, unless otherwise stated. Following 24 hours after mRNA transfection, cells were harvested using Cellstripper^®^ (Corning, catalog# 25-056-Cl) for 15 minutes at 37°C. After cells were lifted they were processed for flow cytometry as described below. For experiments using secreted outputs (i.e. SEAP or secNluc), the supernatant was harvested 24 hours after mRNA transfection for quantification.

### Flow cytometry and data analysis

For experiments using IgG-RELEASE, transfected cells were pelleted at 300 x g for 7 minutes, washed twice using PBS supplemented with 5% BSA, and then incubated with an anti-human IgG antibody conjugated to Alexa Fluor-488 (Southern Biotech Catalog# 2015-30; Lot: C0517-N012B); 1/750 dilution in PBS + 2% BSA) for 1 hour. Following incubation, the cells were washed twice in PBS + 2% BSA and then filtered through a 40 µm sieve. Cells were analyzed by flow cytometry (BioRad ZE5 Cell Analyzer) and Everest Software (version 3.1). FlowJo (version 10.10.0) was used to process the flow cytometry data.

For analysis, cells were gated for cells that were expressing the IgG-RELEASE-T2A-BFP construct (BFP^+^). The median fluorescence intensity (MFI) of AF488 was calculated for wildtype (mCherry^-^) and reporter (mCherry^+^) cells (**Supplementary** Figure 17), and then compared between uninduced and induced conditions. For each condition, an IgG isotype control (Southern Biotech Catalog# 0109-30, Lot: F3218-M690M) was used to correct the MFI of AF488 for non-specific staining.

### Jurkat NIMPLY-CAR T production

Lentivirus for protease components were generated as previously described using 1 well of a 6-well plate of 1.5 ×10^6^ LentiX HEK293 cells transfected plated 24 hrs prior to transfection in 2 mL complete DMEM. Cells were transfected with 0.6 µg pMD2.g, 1.28 µg psPAX2 plasmids, and 1.8 µg of transfer plasmid in 200 µL of JetOptimus buffer and 3 µL of JetOptimus transfection reagent. Lentivirus was harvested 1 day following transfection and concentrated via precipitation, as previously described. Virus was resuspended in 50 uL RPMI. 500 µL of NFAT-GFP reporter line Jurkats were seeded at 250K cells/mL in a 24-well plate and then treated with 50 uL of viral concentrate. Cells were incubated for 72 hours following treatment and were subsequently validated using flow cytometric to quantify the percentage of mCherry^+^ cells and allowed to select under puromycin for 5 days following subsequent transduction. This process was repeated for each protease cell line to transduce a NIMPLY-gated CD19-CAR under either SFFV or EF1*α* promoter control. Cells were then placed under blasticidin selection for 5 days prior to expression validation.

NIMPLY-CAR expression was quantified by primary antibody staining. 5 ×10^4^ cells were isolated by centrifugation at 300xg. Media was removed and cells were washed twice in 150 µL PBS + 2% BSA µL. Cells were then stained in 100 µL of AlexaFluor647 anti-myc antibody (CellSignaling, Catalog# 2233S Lot: 25) at a 1:500 dilution in PBS + 2% BSA for 30 minutes at 4°C. Cells were then washed and resuspended in PBS + 2% BSA and 2 mM EDTA. AlexaFluor647 signal was measured via flow cytometry, where gates were determined by using an untranduced wildtype control.

### Jurkat NIMPLY-CAR T co-culture

NIMPLY-CAR activation co-culture was performed by seeding 5 ×10^4^ NIMPLY-CAR Jurkat cells of each protease condition in 100 uL at a 1:1 ratio of either WT K562 cells or CD19^+^ K562 cells. Cells were incubated at 37°C for 24 hours. Following incubation, cells were spun at 300xg to isolate media for downstream processing, followed by surface staining using BrilliantViolet 421 (BV421) anti-CD69 antibody (BioLegend, Catalog# 310929 Lot: B365637) at a 1:500 dilution. Cells were stained as previously described and analyzed via flow cytometry. NFAT-GFP Jurkat cells were isolated as mCherry^+^/BFP^+^ relative to an untransduced control. Of the mCherry^+^/BFP^+^ cells, GFP and BV421 positive cells were determined relative to an untransduced control as well.

Isolated cell supernatant was stored at 4°C for less than 24 hrs before processing. Secreted protein outputs were quantified using ELISA MAX Deluxe kit for Granzyme B (BioLegend, Catalog # 439207) and Human IL-2 ELISA kit (Abcam, catalog # ab270883) according to manufacturer specifications. Absorbance was measured at 450 nm and 570 nm on a Infinite Pro 200 plate reader (Tecan) and protein concentrations were determined by fitting to a 4-parameter logistic curve.

### Statistical Analysis

For in vitro experiments, values are reported as the average of four biological replicates, and representative of at least two independent biological experiments. For experiments comparing two groups, a Student’s T-test was used to determine statistical significance, following confirmation that equal variance could be assumed (F-test). If equal variance could not be assumed, then a Welch’s correction was used. If the data was not normally distributed, a Mann-Whitney *U* test was used to test for significance. For experiments with more than two groups, a one-way ANOVA with a post hoc Tukey’s test was used for multiple comparisons among the different means. Data was considered statistically significant when the p-value was less than 0.05. Data presented are average +/- SEM, unless otherwise stated. All statistical analysis was performed using Prism 10.0 (GraphPad).

## Acknowledgments

We would like to thank the Bassik lab and Elowitz lab for kindly sharing plasmids used in this work. This work was funded by NIH (R00EB027723, R61CA278398, and DP2OD034951, X.J.G.; NCI F30CA287739-01, A.R.T.), Longevity Impetus grants (X.J.G.), the International Human Frontier Science Program Organization (LT000221/2021-L, A.E.V.), National Science Foundation graduate research fellowships program (DGE-2146755, C.C.C., N.E.), Cellular and Molecular Biology Training Grant (NIH 5 T32 GM007276, C.C.C.), and Stanford Bio-X Bowes Fellowship (A.R.T.).

## Author Contributions

A.E.V., C.C.C., and X.J.G. conceived and directed the study. A.E.V., C.C.C., S.K., and S.Q., performed the experiments validating the RELEASE suite. A.E.V., and N.E. prepared the ARPE-19 cells and in vivo studies. J.K., cultured primary rat cortical neurons for the study. A.R.T., prepared amplicons for next-generation sequencing. A.E.V., C.C.C., and X.J.G. wrote the manuscript. All authors provided feedback on the manuscript.

## Resource Availability

### Lead Contacts

Correspondance

### Data and Code Availability

Raw flow cytometry data is available upon request from the corresponding authors.

### Materials Availability

Plasmids and plasmid maps will be deposited to Addgene.

## References

1. Lim, W.A. (2010). Designing customized cell signalling circuits. Nat. Rev. Mol. Cell Biol. 11, 393–403. 10.1038/nrm2904.

2. Roybal, K.T., and Lim, W.A. (2017). Synthetic immunology: hacking immune cells to expand their therapeutic capabilities. Annu. Rev. Immunol. 35, 229–253. 10.1146/annurev-immunol-051116-052302.

3. Elowitz, M., and Lim, W.A. (2010). Build life to understand it. Nature 468, 889–890. 10.1038/468889a.

4. Andrianantoandro, E., Basu, S., Karig, D.K., and Weiss, R. (2006). Synthetic biology: new engineering rules for an emerging discipline. Mol. Syst. Biol. 2, 2006.0028. 10.1038/msb4100073.

5. Teixeira, A.P., and Fussenegger, M. (2024). Synthetic macromolecular switches for precision control of therapeutic cell functions. Nat. Rev. Bioeng. 10.1038/s44222-024-00235-9.

6. Daringer, N.M., Dudek, R.M., Schwarz, K.A., and Leonard, J.N. (2014). Modular extracellular sensor architecture for engineering mammalian cell-based devices. ACS Synth. Biol. 3, 892–902. 10.1021/sb400128g.

7. Zhang, X., Mille-Fragoso, L.S., Kaseniit, K.E., Lee, A.P., Zhang, M., Call, C.C., Hu, Y., Xie, Y., and Gao, X.J. (2025). Post-transcriptional modular synthetic receptors. Nat. Chem. Biol. 10.1038/s41589-025-01872-w.

8. Morsut, L., Roybal, K.T., Xiong, X., Gordley, R.M., Coyle, S.M., Thomson, M., and Lim, W.A. (2016). Engineering customized cell sensing and response behaviors using synthetic notch receptors. Cell 164, 780–791. 10.1016/j.cell.2016.01.012.

9. Scheller, L., Strittmatter, T., Fuchs, D., Bojar, D., and Fussenegger, M. (2018). Generalized extracellular molecule sensor platform for programming cellular behavior. Nat. Chem. Biol. 14, 723–729. 10.1038/s41589-018-0046-z.

10. Allen, G.M., Frankel, N.W., Reddy, N.R., Bhargava, H.K., Yoshida, M.A., Stark, S.R., Purl, M., Lee, J., Yee, J.L., Yu, W., et al. (2022). Synthetic cytokine circuits that drive T cells into immune-excluded tumors. Science 378, eaba1624. 10.1126/science.aba1624.

11. Ye, H., Xie, M., Xue, S., Charpin-El Hamri, G., Yin, J., Zulewski, H., and Fussenegger, M. (2017). Self-adjusting synthetic gene circuit for correcting insulin resistance. Nat. Biomed. Eng. 1, 0005. 10.1038/s41551-016-0005.

12. Müller, M.M., Arndt, K.M., and Hoffmann, S.A. (2025). Genetic circuits in synthetic biology: broadening the toolbox of regulatory devices. Front. Synth. Biol. 3. 10.3389/fsybi.2025.1548572.

13. Lillacci, G., Benenson, Y., and Khammash, M. (2018). Synthetic control systems for high performance gene expression in mammalian cells. Nucleic Acids Res. 46, 9855–9863. 10.1093/nar/gky795.

14. Nissim, L., Wu, M.-R., Pery, E., Binder-Nissim, A., Suzuki, H.I., Stupp, D., Wehrspaun, C., Tabach, Y., Sharp, P.A., and Lu, T.K. (2017). Synthetic RNA-Based Immunomodulatory Gene Circuits for Cancer Immunotherapy. Cell 171, 1138–1150.e15. 10.1016/j.cell.2017.09.049.

15. Xie, M., and Fussenegger, M. (2018). Designing cell function: assembly of synthetic gene circuits for cell biology applications. Nat. Rev. Mol. Cell Biol. 19, 507–525. 10.1038/s41580-018-0024-z.

16. Stefanov, B.-A., Teixeira, A.P., Mansouri, M., Bertschi, A., Krawczyk, K., Hamri, G.C.-E., Xue, S., and Fussenegger, M. (2021). Genetically encoded protein thermometer enables precise electrothermal control of transgene expression. Adv Sci (Weinh) 8, e2101813. 10.1002/advs.202101813.

17. Das, A.T., Tenenbaum, L., and Berkhout, B. (2016). Tet-On Systems For Doxycycline-inducible Gene Expression. Curr. Gene Ther. 16, 156–167. 10.2174/1566523216666160524144041.

18. Li, H.-S., Israni, D.V., Gagnon, K.A., Gan, K.A., Raymond, M.H., Sander, J.D., Roybal, K.T., Joung, J.K., Wong, W.W., and Khalil, A.S. (2022). Multidimensional control of therapeutic human cell function with synthetic gene circuits. Science 378, 1227–1234. 10.1126/science.ade0156.

19. Gao, Y., Xiong, X., Wong, S., Charles, E.J., Lim, W.A., and Qi, L.S. (2016). Complex transcriptional modulation with orthogonal and inducible dCas9 regulators. Nat. Methods 13, 1043–1049. 10.1038/nmeth.4042.

20. Gao, X.J., Chong, L.S., Kim, M.S., and Elowitz, M.B. (2018). Programmable protein circuits in living cells. Science 361, 1252–1258. 10.1126/science.aat5062.

21. Fink, T., and Jerala, R. (2022). Designed protease-based signaling networks. Curr. Opin. Chem. Biol. 68, 102146. 10.1016/j.cbpa.2022.102146.

22. Vlahos, A.E., Kang, J., Aldrete, C.A., Zhu, R., Chong, L.S., Elowitz, M.B., and Gao, X.J. (2022). Protease-controlled secretion and display of intercellular signals. Nat. Commun. 13, 912. 10.1038/s41467-022-28623-y.

23. Praznik, A., Fink, T., Franko, N., Lonzarić, J., Benčina, M., Jerala, N., Plaper, T., Roškar, S., and Jerala, R. (2022). Regulation of protein secretion through chemical regulation of endoplasmic reticulum retention signal cleavage. Nat. Commun. 13, 1323. 10.1038/s41467-022-28971-9.

24. Mansouri, M., Ray, P.G., Franko, N., Xue, S., and Fussenegger, M. (2023). Design of programmable post-translational switch control platform for on-demand protein secretion in mammalian cells. Nucleic Acids Res. 51, e1. 10.1093/nar/gkac916.

25. Wang, X., Kang, L., Kong, D., Wu, X., Zhou, Y., Yu, G., Dai, D., and Ye, H. (2024). A programmable protease-based protein secretion platform for therapeutic applications. Nat. Chem. Biol. 20, 432–442. 10.1038/s41589-023-01433-z.

26. Aldrete, C.A., An, C., Call, C.C., Gao, X.J., and Vlahos, A.E. (2024). Perspectives on synthetic protein circuits in mammalian cells. Curr. Opin. Biomed. Eng 32. 10.1016/j.cobme.2024.100555.

27. Chen, Z., and Elowitz, M.B. (2021). Programmable protein circuit design. Cell 184, 2284–2301. 10.1016/j.cell.2021.03.007.

28. Mahameed, M., Xue, S., Stefanov, B.-A., Hamri, G.C.-E., and Fussenegger, M. (2022). Engineering a rapid insulin release system controlled by oral drug administration. Adv Sci (Weinh) 9, e2105619. 10.1002/advs.202105619.

29. Fink, T., Lonzarić, J., Praznik, A., Plaper, T., Merljak, E., Leben, K., Jerala, N., Lebar, T., Strmšek, Ž., Lapenta, F., et al. (2019). Design of fast proteolysis-based signaling and logic circuits in mammalian cells. Nat. Chem. Biol. 15, 115–122. 10.1038/s41589-018-0181-6.

30. Zhu, I., Liu, R., Garcia, J.M., Hyrenius-Wittsten, A., Piraner, D.I., Alavi, J., Israni, D.V., Liu, B., Khalil, A.S., and Roybal, K.T. (2022). Modular design of synthetic receptors for programmed gene regulation in cell therapies. Cell 185, 1431–1443.e16. 10.1016/j.cell.2022.03.023.

31. Edelstein, H.I., Donahue, P.S., Muldoon, J.J., Kang, A.K., Dolberg, T.B., Battaglia, L.M., Allchin, E.R., Hong, M., and Leonard, J.N. (2020). Elucidation and refinement of synthetic receptor mechanisms. Synth Biol (Oxf) 5, ysaa017. 10.1093/synbio/ysaa017.

32. Heusser, K., Yuan, H., Neagoe, I., Tarasov, A.I., Ashcroft, F.M., and Schwappach, B. (2006). Scavenging of 14-3-3 proteins reveals their involvement in the cell-surface transport of ATP-sensitive K+ channels. J. Cell Sci. 119, 4353–4363. 10.1242/jcs.03196.

33. Rajan, S., Preisig-Müller, R., Wischmeyer, E., Nehring, R., Hanley, P.J., Renigunta, V., Musset, B., Schlichthörl, G., Derst, C., Karschin, A., et al. (2002). Interaction with 14-3-3 proteins promotes functional expression of the potassium channels TASK-1 and TASK-3. J Physiol (Lond) 545, 13–26. 10.1113/jphysiol.2002.027052.

34. Sharpe, H.J., Stevens, T.J., and Munro, S. (2010). A comprehensive comparison of transmembrane domains reveals organelle-specific properties. Cell 142, 158–169. 10.1016/j.cell.2010.05.037.

35. Munro, S. (1995). An investigation of the role of transmembrane domains in Golgi protein retention. EMBO J. 14, 4695–4704. 10.1002/j.1460-2075.1995.tb00151.x.

36. Jackson, M.R., Nilsson, T., and Peterson, P.A. (1990). Identification of a consensus motif for retention of transmembrane proteins in the endoplasmic reticulum. EMBO J. 9, 3153–3162. 10.1002/j.1460-2075.1990.tb07513.x.

37. Shikano, S., Coblitz, B., Sun, H., and Li, M. (2005). Genetic isolation of transport signals directing cell surface expression. Nat. Cell Biol. 7, 985–992. 10.1038/ncb1297.

38. Liu, Z., Chen, O., Wall, J.B.J., Zheng, M., Zhou, Y., Wang, L., Vaseghi, H.R., Qian, L., and Liu, J. (2017). Systematic comparison of 2A peptides for cloning multi-genes in a polycistronic vector. Sci. Rep. 7, 2193. 10.1038/s41598-017-02460-2.

39. Mrowiec, T., and Schwappach, B. (2006). 14-3-3 proteins in membrane protein transport. Biol. Chem. 387, 1227–1236. 10.1515/BC.2006.152.

40. Okamoto, Y., and Shikano, S. (2012). Phosphorylation-Regulated Cell Surface Expression of Membrane Proteins. In Protein Kinases, G. Da Silva Xavier, ed. (InTech). 10.5772/38031.

41. Shiwarski, D.J., Crilly, S.E., Dates, A., and Puthenveedu, M.A. (2019). Dual RXR motifs regulate nerve growth factor-mediated intracellular retention of the delta opioid receptor. Mol. Biol. Cell 30, 680–690. 10.1091/mbc.E18-05-0292.

42. Shikano, S., and Li, M. (2003). Membrane receptor trafficking: evidence of proximal and distal zones conferred by two independent endoplasmic reticulum localization signals. Proc Natl Acad Sci USA 100, 5783–5788. 10.1073/pnas.1031748100.

43. Wang, B., Yang, H., Liu, Y.C., Jelinek, T., Zhang, L., Ruoslahti, E., and Fu, H. (1999). Isolation of high-affinity peptide antagonists of 14-3-3 proteins by phage display. Biochemistry 38, 12499–12504. 10.1021/bi991353h.

44. Petosa, C., Masters, S.C., Bankston, L.A., Pohl, J., Wang, B., Fu, H., and Liddington, R.C. (1998). 14-3-3zeta binds a phosphorylated Raf peptide and an unphosphorylated peptide via its conserved amphipathic groove. J. Biol. Chem. 273, 16305–16310. 10.1074/jbc.273.26.16305.

45. McFarlane, A., Pohler, E., and Moraga, I. (2023). Molecular and cellular factors determining the functional pleiotropy of cytokines. FEBS J. 290, 2525–2552. 10.1111/febs.16420.

46. Martinez-Sanchez, M.E., Huerta, L., Alvarez-Buylla, E.R., and Villarreal Luján, C. (2018). Role of cytokine combinations on CD4+ T cell differentiation, partial polarization, and plasticity: continuous network modeling approach. Front. Physiol. 9, 877. 10.3389/fphys.2018.00877.

47. Biernacka, A., Dobaczewski, M., and Frangogiannis, N.G. (2011). TGF-β signaling in fibrosis. Growth Factors 29, 196–202. 10.3109/08977194.2011.595714.

48. Zhang, F., Cong, L., Lodato, S., Kosuri, S., Church, G.M., and Arlotta, P. (2011). Efficient construction of sequence-specific TAL effectors for modulating mammalian transcription. Nat. Biotechnol. 29, 149–153. 10.1038/nbt.1775.

49. Soto, L.F., Li, Z., Santoso, C.S., Berenson, A., Ho, I., Shen, V.X., Yuan, S., and Fuxman Bass, J.I. (2022). Compendium of human transcription factor effector domains. Mol. Cell 82, 514–526. 10.1016/j.molcel.2021.11.007.

50. Shahein, A., López-Malo, M., Istomin, I., Olson, E.J., Cheng, S., and Maerkl, S.J. (2022). Systematic analysis of low-affinity transcription factor binding site clusters in vitro and in vivo establishes their functional relevance. Nat. Commun. 13, 5273. 10.1038/s41467-022-32971-0.

51. Kapust, R.B., Tözsér, J., Copeland, T.D., and Waugh, D.S. (2002). The P1’ specificity of tobacco etch virus protease. Biochem. Biophys. Res. Commun. 294, 949–955. 10.1016/S0006-291X(02)00574-0.

52. Wang, W., Wildes, C.P., Pattarabanjird, T., Sanchez, M.I., Glober, G.F., Matthews, G.A., Tye, K.M., and Ting, A.Y. (2017). A light- and calcium-gated transcription factor for imaging and manipulating activated neurons. Nat. Biotechnol. 35, 864–871. 10.1038/nbt.3909.

53. Park, J.W., Reed, J.R., Brignac-Huber, L.M., and Backes, W.L. (2014). Cytochrome P450 system proteins reside in different regions of the endoplasmic reticulum. Biochem. J. 464, 241–249. 10.1042/BJ20140787.

54. Szczesna-Skorupa, E., and Kemper, B. (2000). Endoplasmic reticulum retention determinants in the transmembrane and linker domains of cytochrome P450 2C1. J. Biol. Chem. 275, 19409–19415. 10.1074/jbc.M002394200.

55. Verbič, A., Lebar, T., Praznik, A., and Jerala, R. (2024). Subunits of an E3 ligase complex as degrons for efficient degradation of cytosolic, nuclear, and membrane proteins. ACS Synth. Biol. 13, 792–803. 10.1021/acssynbio.3c00588.

56. Fernandez-Rodriguez, J., and Voigt, C.A. (2016). Post-translational control of genetic circuits using Potyvirus proteases. Nucleic Acids Res. 44, 6493–6502. 10.1093/nar/gkw537.

57. Lee, J.M., Hammarén, H.M., Savitski, M.M., and Baek, S.H. (2023). Control of protein stability by post-translational modifications. Nat. Commun. 14, 201. 10.1038/s41467-023-35795-8.

58. White, S.H., and von Heijne, G. (2005). Transmembrane helices before, during, and after insertion. Curr. Opin. Struct. Biol. 15, 378–386. 10.1016/j.sbi.2005.07.004.

59. Alabanza, L., Pegues, M., Geldres, C., Shi, V., Wiltzius, J.J.W., Sievers, S.A., Yang, S., and Kochenderfer, J.N. (2017). Function of Novel Anti-CD19 Chimeric Antigen Receptors with Human Variable Regions Is Affected by Hinge and Transmembrane Domains. Mol. Ther. 25, 2452–2465. 10.1016/j.ymthe.2017.07.013.

60. Tycko, J., DelRosso, N., Hess, G.T., Aradhana, Banerjee, A., Mukund, A., Van, M.V., Ego, B.K., Yao, D., Spees, K., et al. (2020). High-Throughput Discovery and Characterization of Human Transcriptional Effectors. Cell 183, 2020–2035.e16. 10.1016/j.cell.2020.11.024.

61. Bulcha, J.T., Wang, Y., Ma, H., Tai, P.W.L., and Gao, G. (2021). Viral vector platforms within the gene therapy landscape. Signal Transduct. Target. Ther. 6, 53. 10.1038/s41392-021-00487-6.

62. Wu, Z., Yang, H., and Colosi, P. (2010). Effect of genome size on AAV vector packaging. Mol. Ther. 18, 80–86. 10.1038/mt.2009.255.

63. Kalidasan, V., Ng, W.H., Ishola, O.A., Ravichantar, N., Tan, J.J., and Das, K.T. (2021). A guide in lentiviral vector production for hard-to-transfect cells, using cardiac-derived c-kit expressing cells as a model system. Sci. Rep. 11, 19265. 10.1038/s41598-021-98657-7.

64. Kumar, M., Keller, B., Makalou, N., and Sutton, R.E. (2001). Systematic determination of the packaging limit of lentiviral vectors. Hum. Gene Ther. 12, 1893–1905. 10.1089/104303401753153947.

65. Szymczak, A.L., Workman, C.J., Wang, Y., Vignali, K.M., Dilioglou, S., Vanin, E.F., and Vignali, D.A.A. (2004). Correction of multi-gene deficiency in vivo using a single “self-cleaving” 2A peptide-based retroviral vector. Nat. Biotechnol. 22, 589–594. 10.1038/nbt957.

66. Juillerat, A., Tkach, D., Busser, B.W., Temburni, S., Valton, J., Duclert, A., Poirot, L., Depil, S., and Duchateau, P. (2019). Modulation of chimeric antigen receptor surface expression by a small molecule switch. BMC Biotechnol. 19, 44. 10.1186/s12896-019-0537-3.

67. Chung, H.K., and Lin, M.Z. (2020). On the cutting edge: protease-based methods for sensing and controlling cell biology. Nat. Methods 17, 885–896. 10.1038/s41592-020-0891-z.

68. Kanuga, N., Winton, H.L., Beauchéne, L., Koman, A., Zerbib, A., Halford, S., Couraud, P.-O., Keegan, D., Coffey, P., Lund, R.D., et al. (2002). Characterization of Genetically Modified Human Retinal Pigment Epithelial Cells Developed for In Vitro and Transplantation Studies | IOVS | ARVO Journals. Investigative Ophthalmology & Visual Science.

69. Nash, A.M., Jarvis, M.I., Aghlara-Fotovat, S., Mukherjee, S., Hernandez, A., Hecht, A.D., Rios, P.D., Ghani, S., Joshi, I., Isa, D., et al. (2022). Clinically translatable cytokine delivery platform for eradication of intraperitoneal tumors. Sci. Adv. 8, eabm1032. 10.1126/sciadv.abm1032.

70. Somanathan, S., Breous, E., Bell, P., and Wilson, J.M. (2010). AAV vectors avoid inflammatory signals necessary to render transduced hepatocyte targets for destructive T cells. Mol. Ther. 18, 977–982. 10.1038/mt.2010.40.

71. Wang, J.-H., Gessler, D.J., Zhan, W., Gallagher, T.L., and Gao, G. (2024). Adeno-associated virus as a delivery vector for gene therapy of human diseases. Signal Transduct. Target. Ther. 9, 78. 10.1038/s41392-024-01780-w.

72. Choi, J.-H., Yu, N.-K., Baek, G.-C., Bakes, J., Seo, D., Nam, H.J., Baek, S.H., Lim, C.-S., Lee, Y.-S., and Kaang, B.-K. (2014). Optimization of AAV expression cassettes to improve packaging capacity and transgene expression in neurons. Mol. Brain 7, 17. 10.1186/1756-6606-7-17.

73. Aldrete, C.A., Call, C.C., Sant’Anna, L.E., Vlahos, A.E., Pei, J., Cong, Q., and Gao, X.J. (2025). Orthogonalized human protease control of secreted signals. Nat. Chem. Biol. 10.1038/s41589-024-01831-x.

74. Lu, A., Moeller, L., Moore, S., Xia, S., Ho, K., Zhang, E., Budde, M.W., Larson, H., Ahmed Diaz, A., Gu, B., et al. (2025). Engineered protein circuits for cancer therapy. BioRxiv. 10.1101/2025.04.16.647665.

75. Kowalski, P.S., Rudra, A., Miao, L., and Anderson, D.G. (2019). Delivering the Messenger: Advances in Technologies for Therapeutic mRNA Delivery. Mol. Ther. 27, 710–728. 10.1016/j.ymthe.2019.02.012.

76. Parayath, N.N., Stephan, S.B., Koehne, A.L., Nelson, P.S., and Stephan, M.T. (2020). In vitro-transcribed antigen receptor mRNA nanocarriers for transient expression in circulating T cells in vivo. Nat. Commun. 11, 6080. 10.1038/s41467-020-19486-2.

77. Cappell, K.M., and Kochenderfer, J.N. (2023). Long-term outcomes following CAR T cell therapy: what we know so far. Nat. Rev. Clin. Oncol. 20, 359–371. 10.1038/s41571-023-00754-1.

78. Flugel, C.L., Majzner, R.G., Krenciute, G., Dotti, G., Riddell, S.R., Wagner, D.L., and Abou-El-Enein, M. (2023). Overcoming on-target, off-tumour toxicity of CAR T cell therapy for solid tumours. Nat. Rev. Clin. Oncol. 20, 49–62. 10.1038/s41571-022-00704-3.

79. Wu, J., Wu, W., Zhou, B., and Li, B. (2024). Chimeric antigen receptor therapy meets mRNA technology. Trends Biotechnol. 42, 228–240. 10.1016/j.tibtech.2023.08.005.

80. Finn, P., Chavez, M., Chen, X., Wang, H., Rane, D.A., Gurjar, J., and Qi, L.S. (2023). Drug-Mediated Control of Receptor Valency Enhances Immune Cell Potency. BioRxiv. 10.1101/2023.01.04.522664.

81. Cibrián, D., and Sánchez-Madrid, F. (2017). CD69: from activation marker to metabolic gatekeeper. Eur. J. Immunol. 47, 946–953. 10.1002/eji.201646837.

82. Wang, W., and Shakes, D.C. (1996). Molecular evolution of the 14-3-3 protein family. J. Mol. Evol. 43, 384–398. 10.1007/BF02339012.

83. Zhou, J., Ge, Q., Wang, D., Guo, Q., and Tao, Y. (2023). Engineering a modular double-transmembrane synthetic receptor system for customizing cellular programs. Cell Rep. 42, 112385. 10.1016/j.celrep.2023.112385.

84. Cabrera, A., Edelstein, H.I., Glykofrydis, F., Love, K.S., Palacios, S., Tycko, J., Zhang, M., Lensch, S., Shields, C.E., Livingston, M., et al. (2022). The sound of silence: Transgene silencing in mammalian cell engineering. Cell Syst. 13, 950–973. 10.1016/j.cels.2022.11.005.

85. Zimak, J., Wagoner, Z.W., Nelson, N., Waechtler, B., Schlosser, H., Kopecky, M., Wu, J., and Zhao, W. (2021). Epigenetic silencing directs expression heterogeneity of stably integrated multi-transcript unit genetic circuits. Sci. Rep. 11, 2424. 10.1038/s41598-021-81975-1.

86. Jüttner, J., Szabo, A., Gross-Scherf, B., Morikawa, R.K., Rompani, S.B., Hantz, P., Szikra, T., Esposti, F., Cowan, C.S., Bharioke, A., et al. (2019). Targeting neuronal and glial cell types with synthetic promoter AAVs in mice, non-human primates and humans. Nat. Neurosci. 22, 1345–1356. 10.1038/s41593-019-0431-2.

87. Taskiran, I.I., Spanier, K.I., Dickmänken, H., Kempynck, N., Pančíková, A., Ekşi, E.C., Hulselmans, G., Ismail, J.N., Theunis, K., Vandepoel, R., et al. (2024). Cell-type-directed design of synthetic enhancers. Nature 626, 212–220. 10.1038/s41586-023-06936-2.

88. Wu, M.-R., Nissim, L., Stupp, D., Pery, E., Binder-Nissim, A., Weisinger, K., Enghuus, C., Palacios, S.R., Humphrey, M., Zhang, Z., et al. (2019). A high-throughput screening and computation platform for identifying synthetic promoters with enhanced cell-state specificity (SPECS). Nat. Commun. 10, 2880. 10.1038/s41467-019-10912-8.

89. Rivera, V.M., Berk, L., and Clackson, T. (2012). Dimerizer-mediated regulation of gene expression in vivo. Cold Spring Harb. Protoc. 2012, 821–824. 10.1101/pdb.prot070144.

90. Su, Y., Walker, J.R., Park, Y., Smith, T.P., Liu, L.X., Hall, M.P., Labanieh, L., Hurst, R., Wang, D.C., Encell, L.P., et al. (2020). Novel NanoLuc substrates enable bright two-population bioluminescence imaging in animals. Nat. Methods 17, 852–860. 10.1038/s41592-020-0889-6.

91. UniProt Consortium (2023). Uniprot: the universal protein knowledgebase in 2023. Nucleic Acids Res. 51, D523–D531. 10.1093/nar/gkac1052.

92. Zulkower, V., and Rosser, S. (2020). DNA Chisel, a versatile sequence optimizer. Bioinformatics 36, 4508–4509. 10.1093/bioinformatics/btaa558.

93. Fang, E., Liu, X., Li, M., Zhang, Z., Song, L., Zhu, B., Wu, X., Liu, J., Zhao, D., and Li, Y. (2022). Advances in COVID-19 mRNA vaccine development. Signal Transduct. Target. Ther. 7, 94. 10.1038/s41392-022-00950-y.

