## Supplementary Figures for "A complete suite for compact programming of protein secretion"

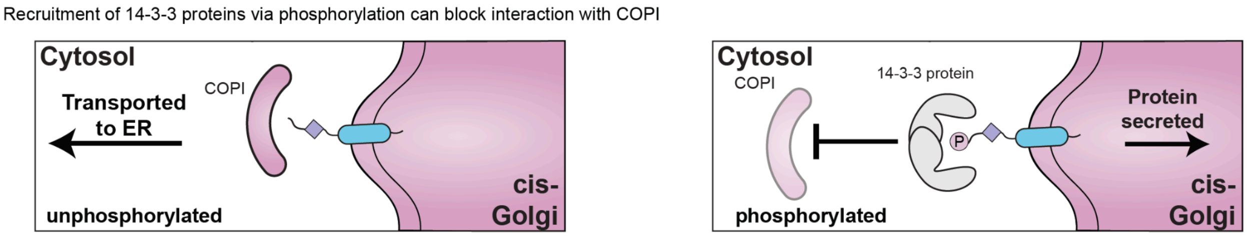


**Supplementary Figure 1:** A schematic of phosphorylation-dependent recruitment of endogenous 14-3-3 scaffolding proteins to control the surface expression of certain membrane proteins. Recruitment of 14-3-3 proteins can block the recognition of ER-retention motifs from the COPI retrograde transport complex.

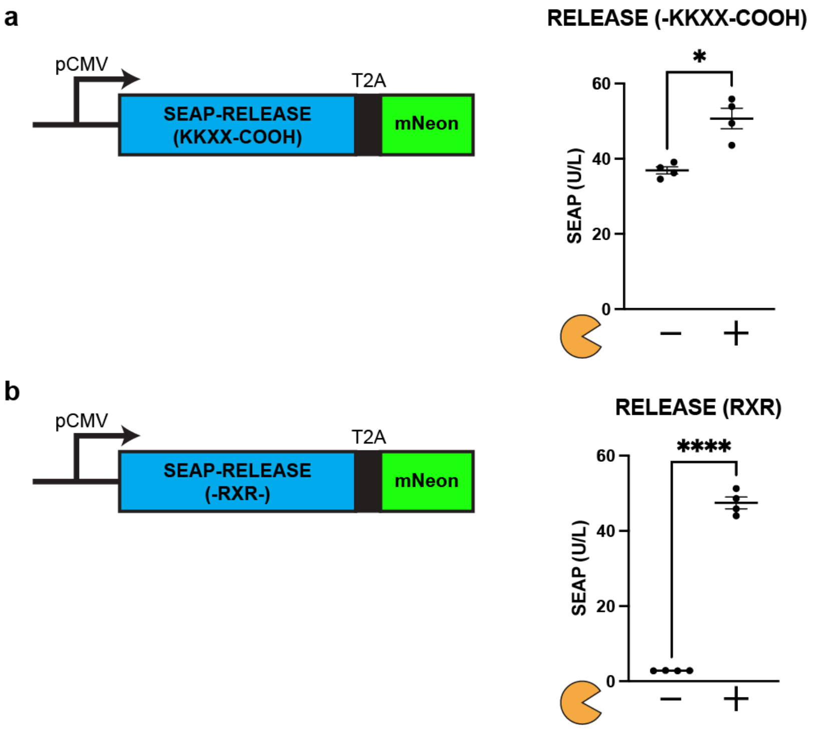


**Supplementary Figure 2: a)** The 2A self-cleaving peptides leave a scar at the C-terminus, and the original RELEASE platform using the dilysine motif loses its ER-retention capabilities, since it must be terminal. **b)** In comparison, RELEASE variants using the diarginine motif (-RXR-) preserved retention capabilities. Mean values were calculated from four replications. The error bars represent ± SEM. The results are representative of at least two independent experiments; significance was tested using an unpaired two-tailed Student's *t*-test between the two indicated conditions for each experiment. *$p < 0.05$, ****$p < 0.001$.

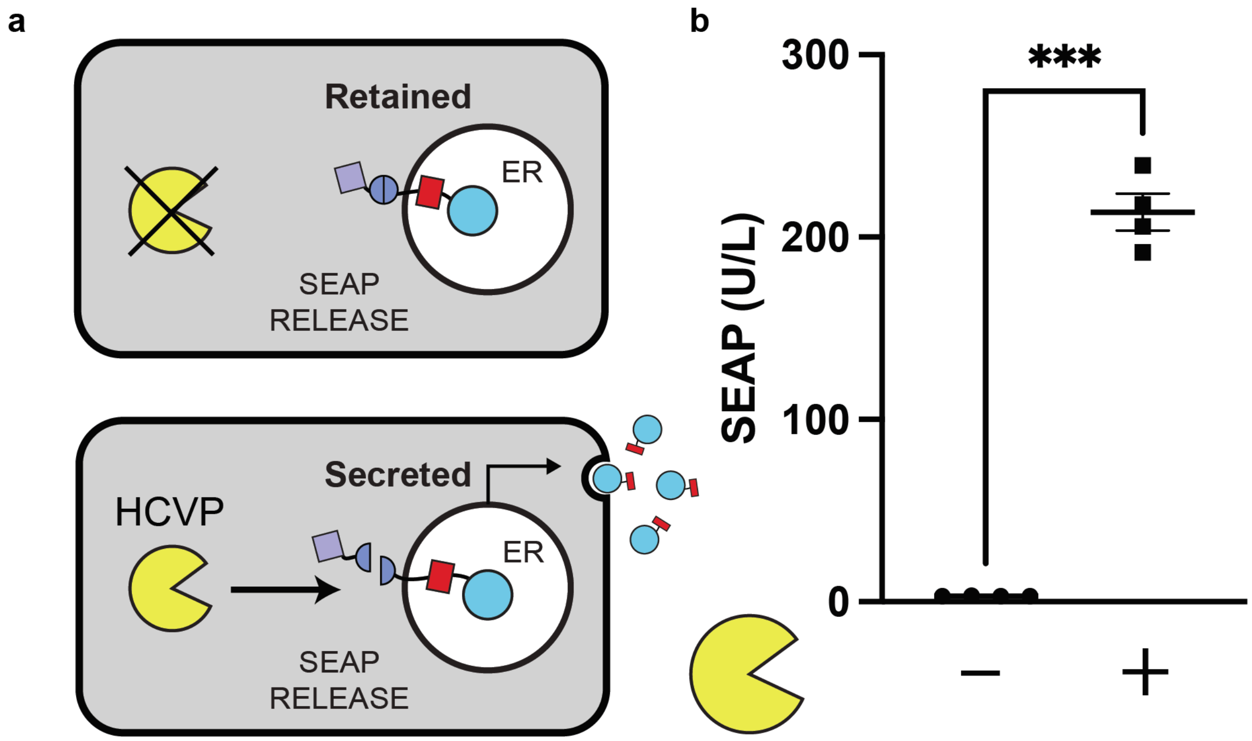


**<u>Supplementary Figure 3:</u> a)** Schematic representation of HCVP-inducible RELEASE for controlling protein secretion. **b)** Switching the cut site of RELEASE to be compatible with HCVP resulted in a significant increase in SEAP secretion when co-expressed with HCVP. Each dot represents a biological replicate. Mean values were calculated from four biological replicates. The error bars represent ± SEM. The results are representative of at least two independent experiments; significance was tested using an unpaired two-tailed Student's *t*-test between the two indicated conditions. ****$p < 0.0001$.

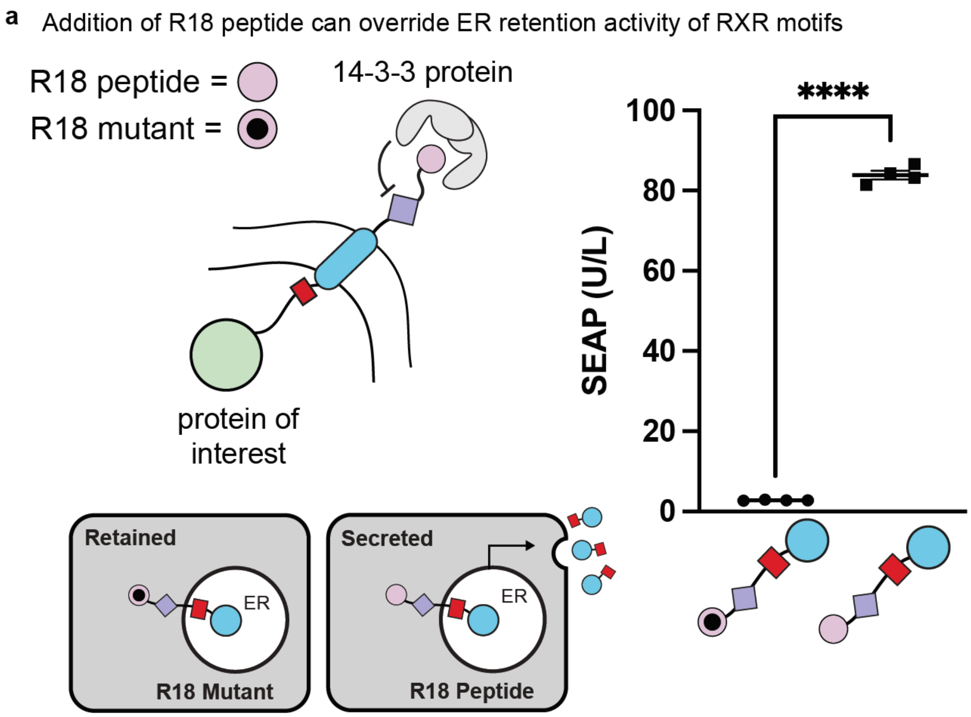


**Supplementary Figure 4:** The fusion of the 14-3-3 antagonist (R18 peptide) to the C-terminus of RELEASE inhibited ER retention activity and resulted in the secretion of SEAP. In comparison, a fused R18 mutant which had two acidic residues required for 14-3-3 recruitment mutated into lysines (D12K and E14K) could not overcome the retention capabilities of RELEASE, and SEAP was not secreted. Each dot represents a biological replicate. Mean values were calculated from four replicates. The error bars represent ± SEM. The results are representative of at least two independent experiments; significance was tested using an unpaired two-tailed Student's *t*-test between the two indicated conditions for each experiment. ****$p < 0.0001$.

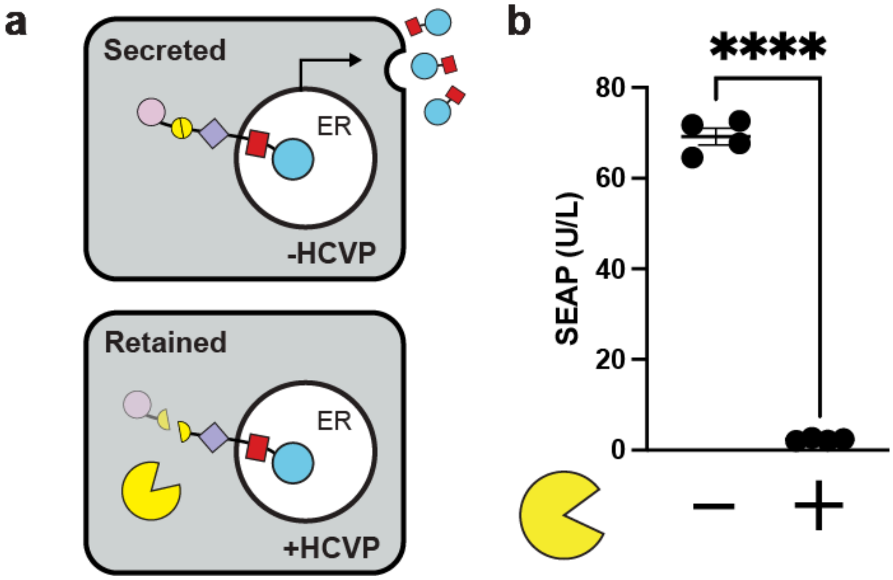


**<u>Supplementary Figure 5:</u> a)** Schematic of HCVP-inducible RELEASE-NOT for the negative regulation of protein secretion. **b)** Similar to RELEASE, by switching the cytoplasmic protease cut site of RELEASE-NOT, additional variants responsive to different proteases such as HCVP can be engineered. In the absence of HCVP, SEAP is secreted. Co-expression of HCVP resulted in a significant decrease in the secretion of SEAP. Mean values were calculated from four replicates. The error bars represent ± SEM. The results are representative of at least two independent experiments; significance was tested using an unpaired two-tailed Student's *t*-test between the two indicated conditions for each experiment. $^{****}p < 0.0001$.

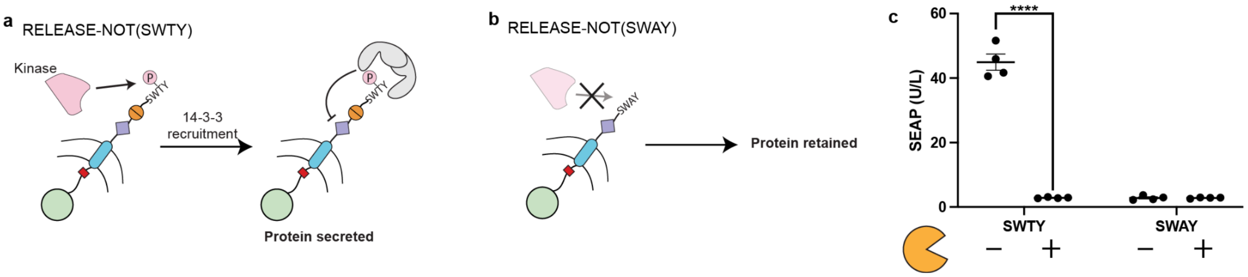


**<u>Supplementary Figure 6:</u>** a) Schematic of RELEASE-NOT using the SWTY motif to enable phosphorylation-dependent recruitment of 14-3-3 proteins to block the retention activity of RXR. **b)** Schematic of RELEASE-NOT where the phospho-responsive threonine residue is mutated to alanine (SWTY-COOH → SWAY-COOH). **c)** When co-expressing TEVP we observed a significant decrease in SEAP secretion only when using the RELEASE-NOT construct with the SWTY motif. RELEASE-NOT with the SWAY motif did not result in any secretion of SEAP. Each dot represents a biological replicate. Mean values were calculated from four replicates. The error bars represent ± SEM. The results are representative of at least two independent experiments; significance was tested using a one-way ANOVA with a Tukey's post-hoc comparison test was used to assess significance. $^{****}p < 0.0001$.

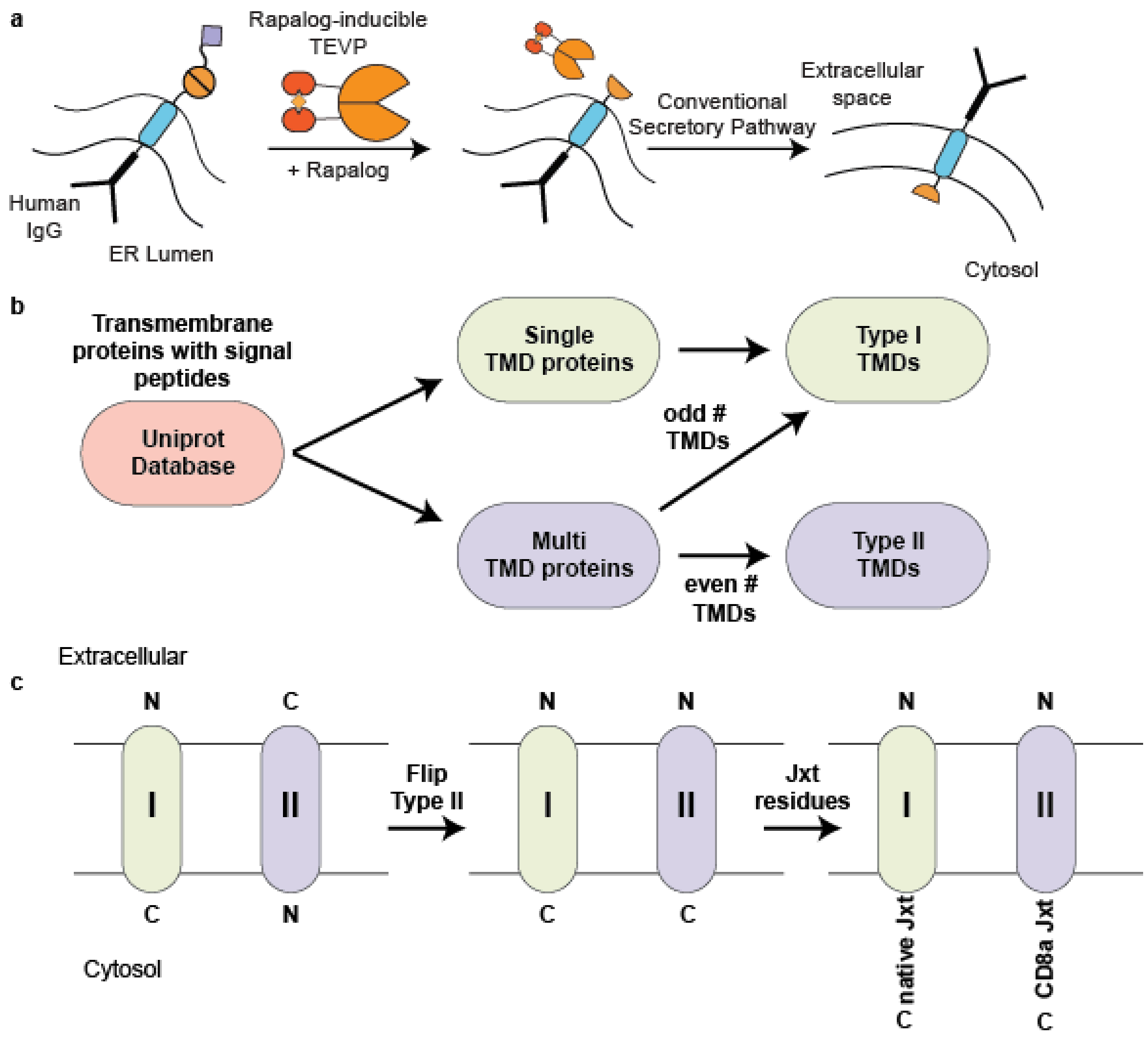


**<u>Supplementary Figure 7:</u> a)** Schematic representation of IgG-RELEASE. **b)** Schematic representation of the transmembrane domain (TMD) library design. First, proteins containing transmembrane domains and signal peptides were queried from the UNIPROT database. Proteins containing a single TMD were already in a Type I orientation, compatible with RELEASE. Proteins containing multi-spanning TMDs had each TMD separated depending on whether they originally adopted a Type I or Type II orientation. The first TMD for each of these multi-spanning TMD proteins has a Type I orientation due to the signal peptide, followed by a TMD with a Type II orientation, alternating between the two orientations. **c)** Within the library, Type I TMDs with their native juxtamembrane (jxt) residues were also included as a separate element. To ensure compatibility with RELEASE, Type II TMDs had their sequences flipped and included within the library. We kept the naming convention the same. For each of these Type II TMDs, we appended the jxt residues from CD8α.

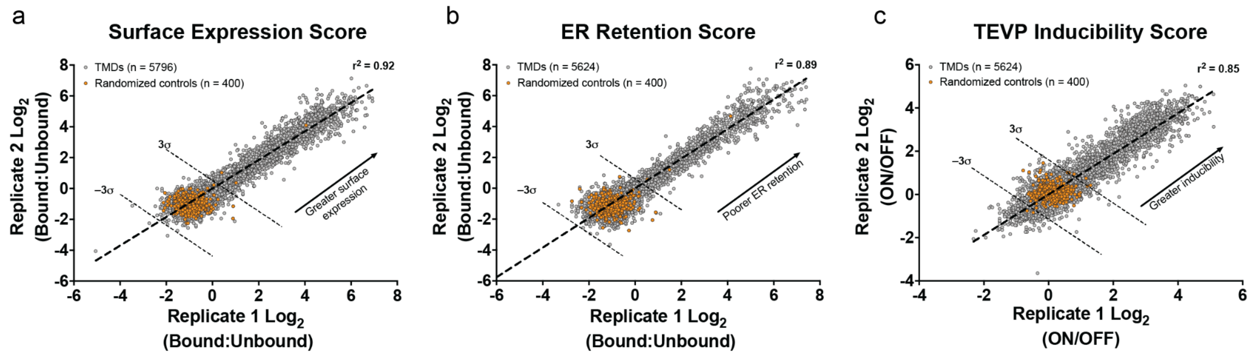


**Supplementary Figure 8: a)** Reproducibility from two biological replicates for surface expression score. **b)** Reproducibility from two biological replicates for ER retention score. **c)** Reproducibility from two biological replicates for TEVP inducibility score. For each graph, the randomized controls, representing the negative population are coloured orange.

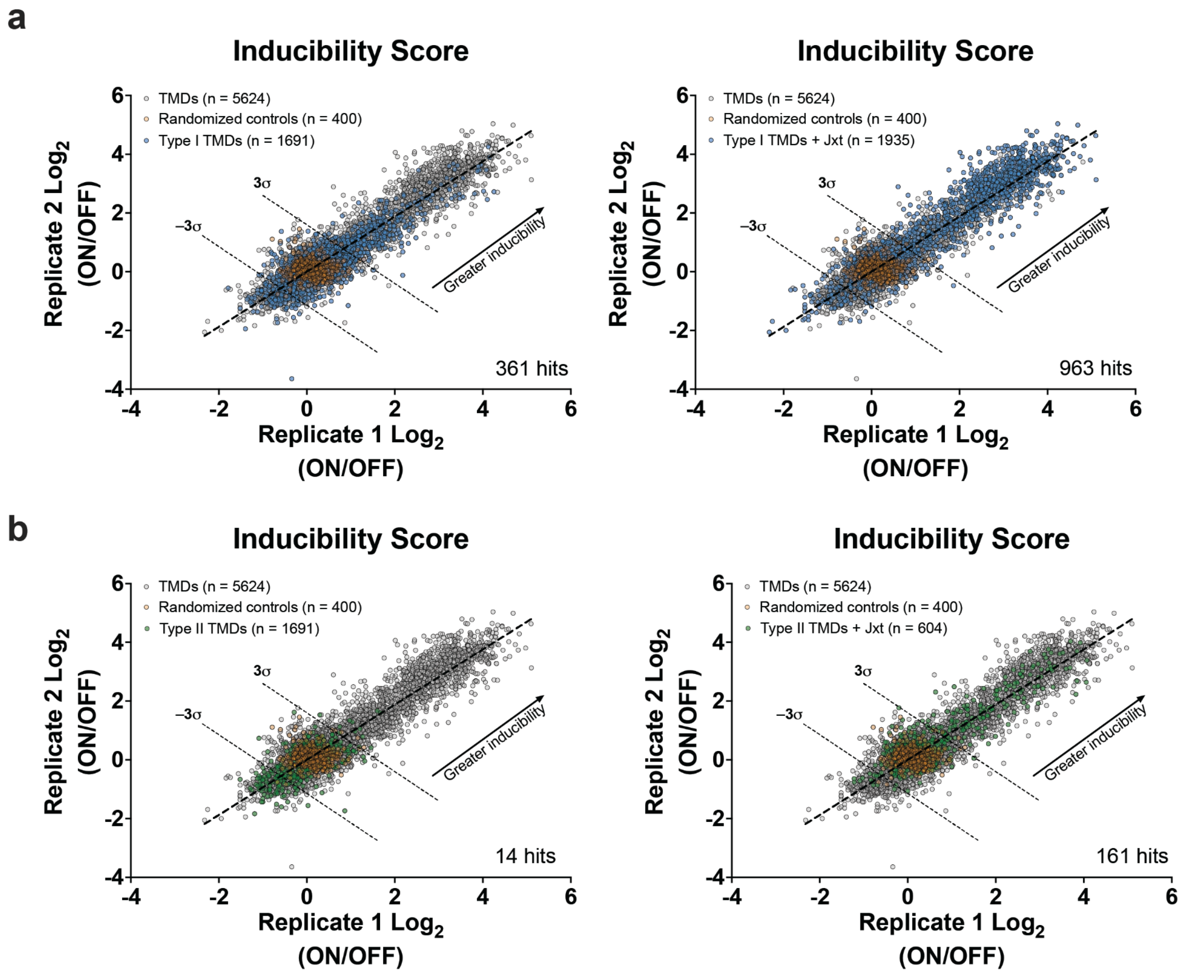


**Supplementary Figure 9: a)** Inducibility scores for RELEASE variants using type I TMDs with and without juxtamembrane residues, which were coloured blue. The hit threshold is set to 3 SDs above the mean of the negative controls, coloured orange. **b)** Inducibility scores for

RELEASE variants using type II TMDs with and without juxtamembrane residues, which were coloured green. The hit threshold is set to 3 SDs above the mean of the negative controls, coloured orange.

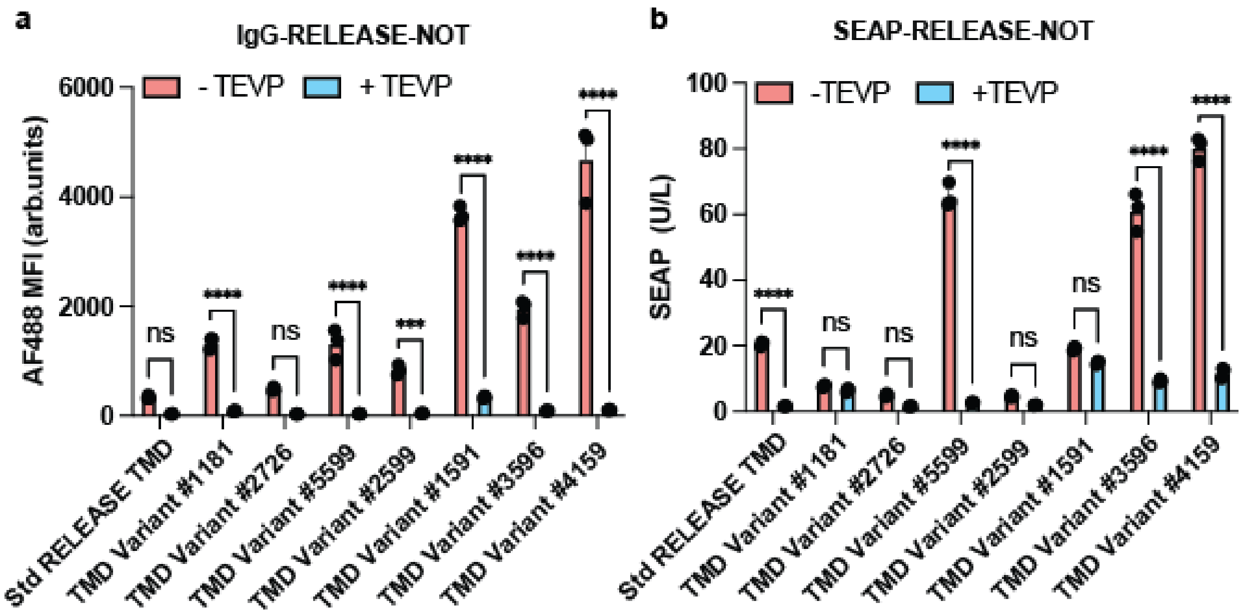


**Supplementary Figure 10:** **a)** Bar plots of the surface expression of IgG-RELEASE-NOT variants with different TMDs, co-expressed with and without TEVP. **b)** Bar plots of SEAP secretion of SEAP-RELEASE-NOT variants with TMDs, co-expressed with and without TEVP. For both plots, RELEASE-NOT variants were ordered based on ascending library TEVP induction scores. ***$p$ <0.001, ****$p$ <0.0001.

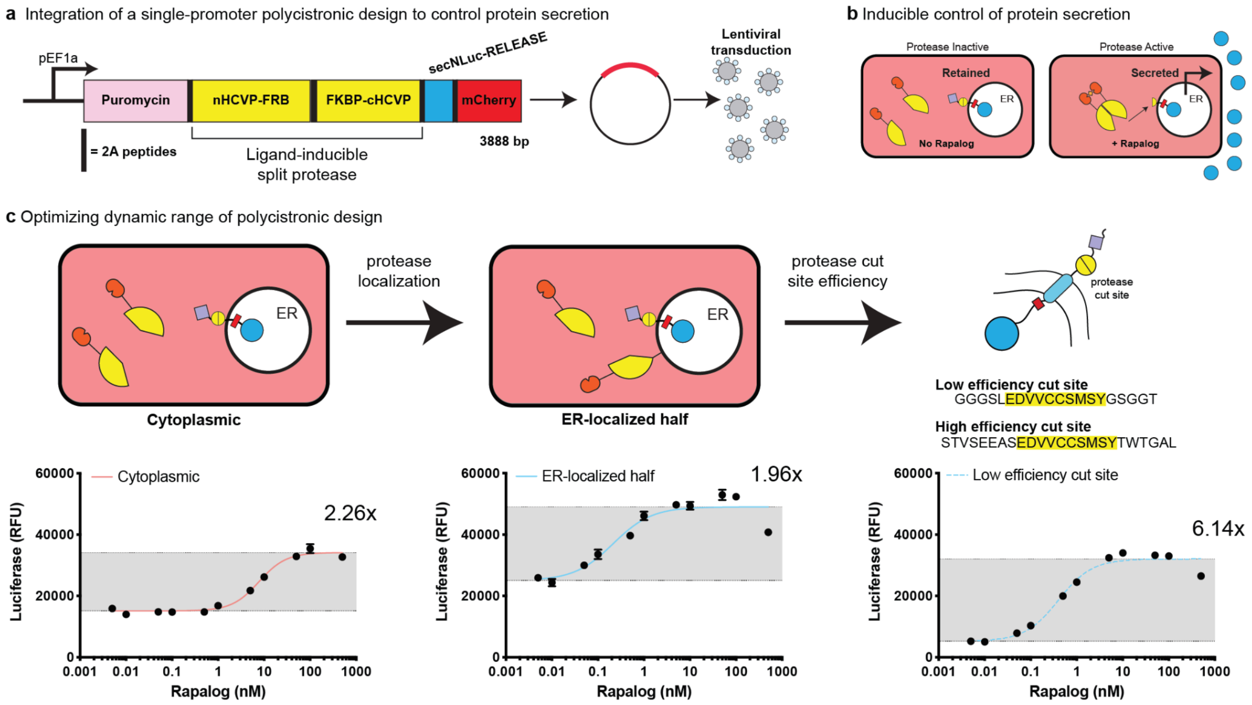


**Supplementary Figure 11:** **a)** Schematic of the polycistronic gene encoding all the protein components required to stably engineer mammalian cells for inducible protein secretion via lentiviral transduction. **b)** To control the secretion of the secNluc reporter, a split HCVP protease fused to the FKBP and FRB dimerization domains were expressed in the cytoplasm. Only in the presence of rapalog would the split protease be reconstituted and induce protein secretion. **c)** Input-output curves for engineered HEK293 cells where secNluc functions as the output and rapalog concentration serves as the input. The curve represents a Hill equation with a variable slope, and the grey box represents the top and bottom plateaus of the fit. To increase Nluc secretion, one half of the split HCVP was localized to the ER via the p450-motif, however this design resulted in an increase in background secretion and a reduction in the dynamic range. To reduce the background and increase the dynamic range, the cleavage efficiency of the HCVP cut site of RELEASE was reduced by modifying the flanking residues, as previously described[1]. Mean values were validated from four replicates. The error bars represent +/- SEM. The results are representative of at least two independent experiments.

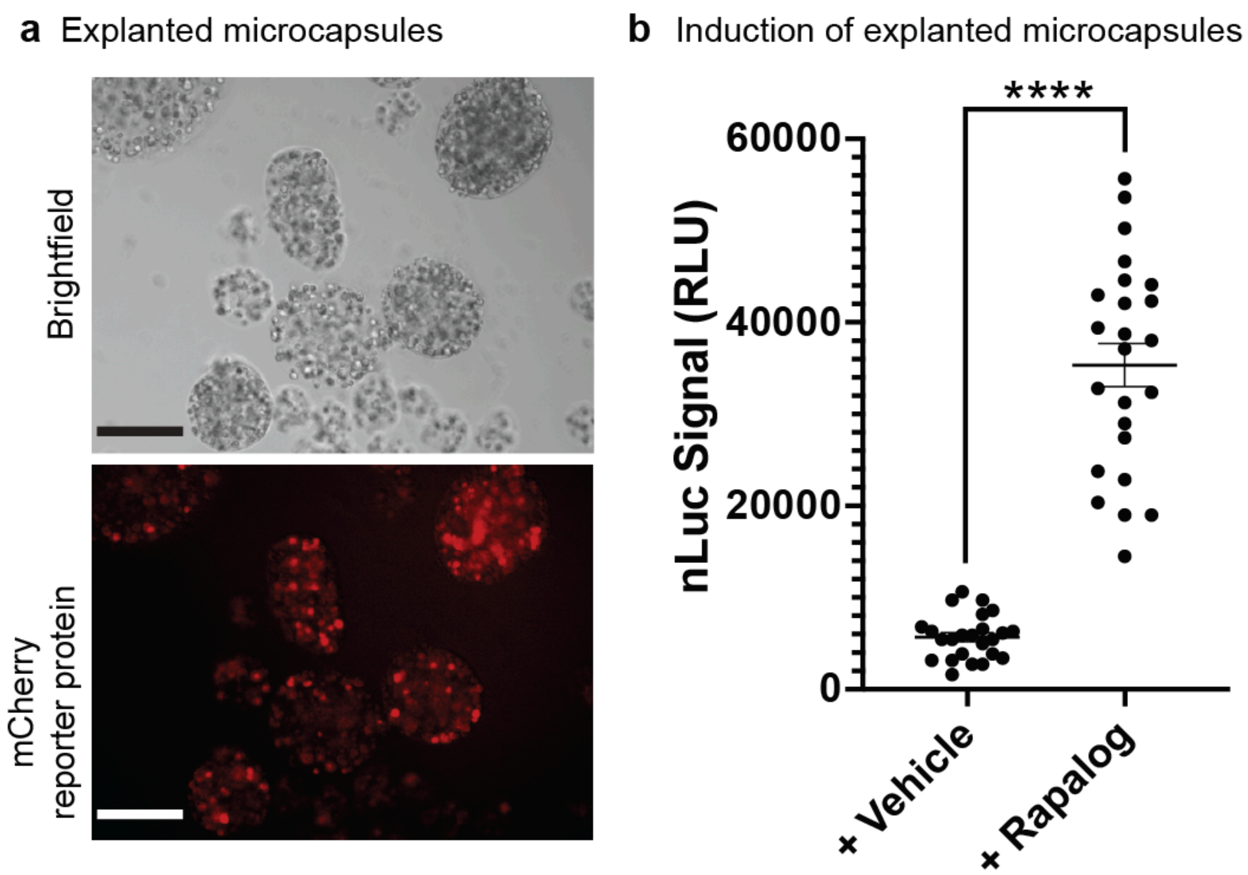


**Supplementary Figure 12:** **a)** Representative microscopy images of explanted alginate microcapsules containing engineered ARPE-19 cells. Scale bars = 300 µm. **b)** Dot plot comparing rapalog-inducible secNluc secretion of explanted microcapsules with engineered ARPE-19 cells. Each dot represents a biological replicate made of 25 microcapsules. Significance was tested using an unpaired two-tailed Student's *t*-test between the two conditions. ****$p < 0.0001$.

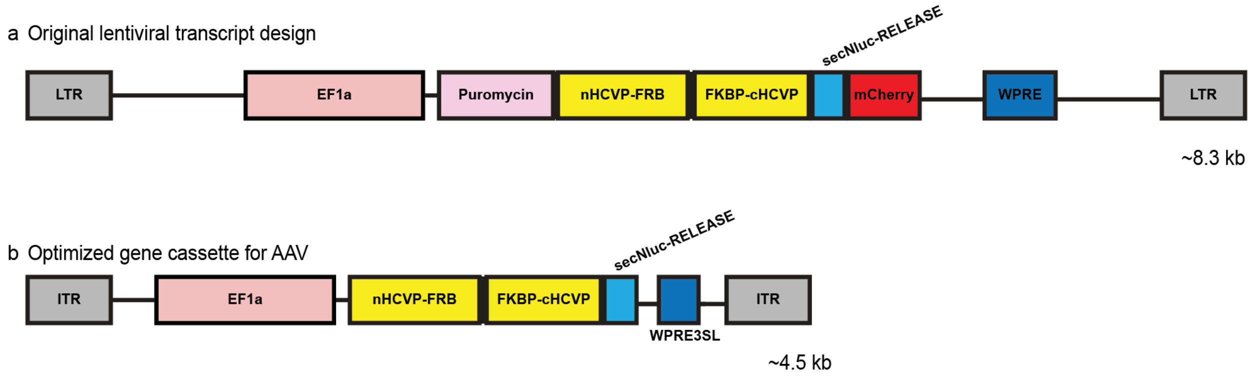


**Supplementary Figure 13:** **a)** Schematic representation of lentiviral transfer plasmid encoding the puromycin resistance gene, a rapalog-inducible HCVP, secNluc-RELEASE and the mCherry reporter protein. **b)** Schematic representation of AAV transfer plasmid encoding a synthetic protein circuit for rapalog-inducible control of secNluc via RELEASE.

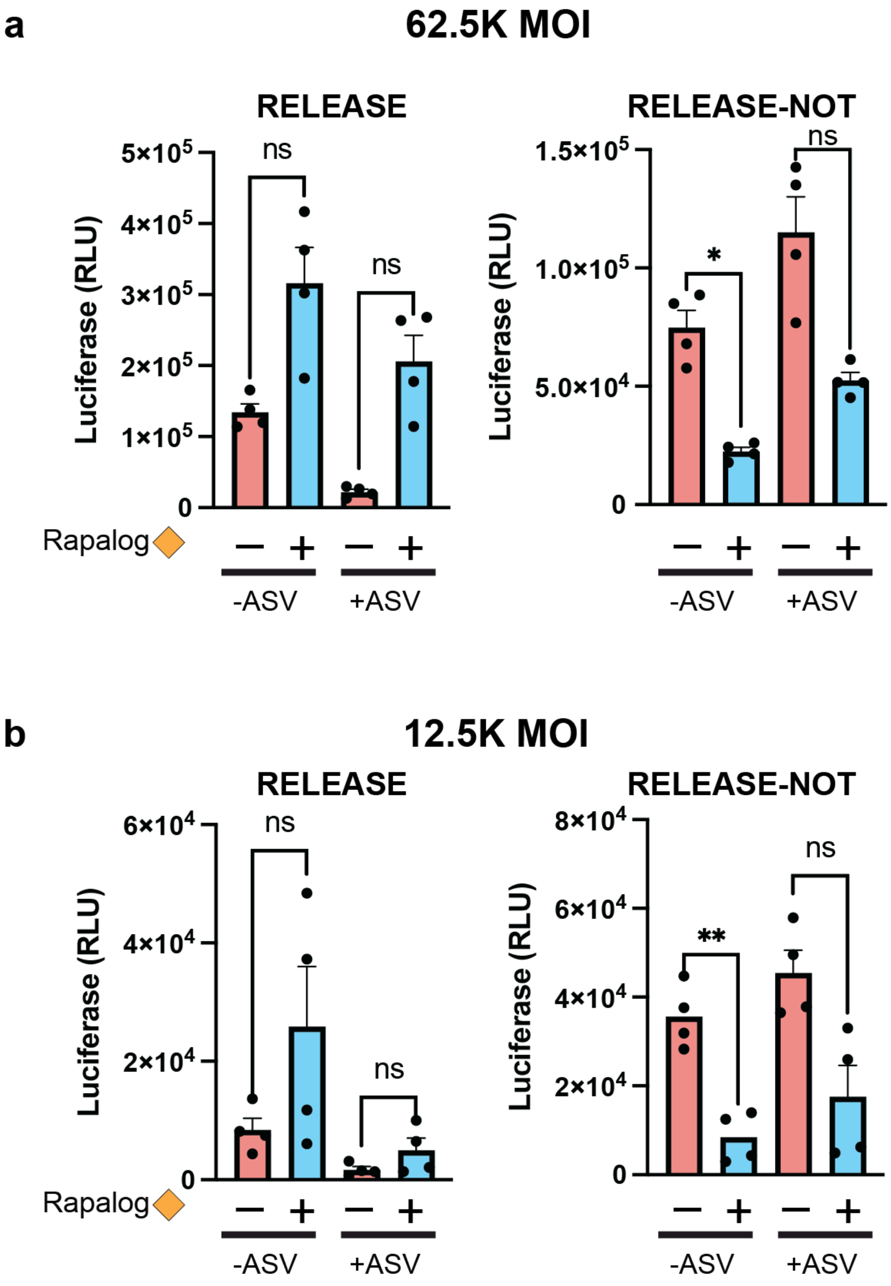


**Supplementary Figure 14:** **a)** Dot plots of secreted nanoLuc values from primary rat cortical neurons transduced with 62.5K MOI of AAV encoding RELEASE or RELEASE-NOT. Primary rat cortical neurons were incubated with and without rapalog, or asunaprevir (ASV). **b)** Dot plots from primary rat cortical neurons transduced with 12.5K MOI of AAV encoding RELEASE or RELEASE-NOT. The absolute amount of secreted nanoLuc was dependent on the MOI used for the AAV transduction. N = 4 biological replicates. Statistical significance was calculated using a two-way ANOVA with Bonferroni's multiple comparisons test. *$p < 0.05$; **$p < 0.01$.

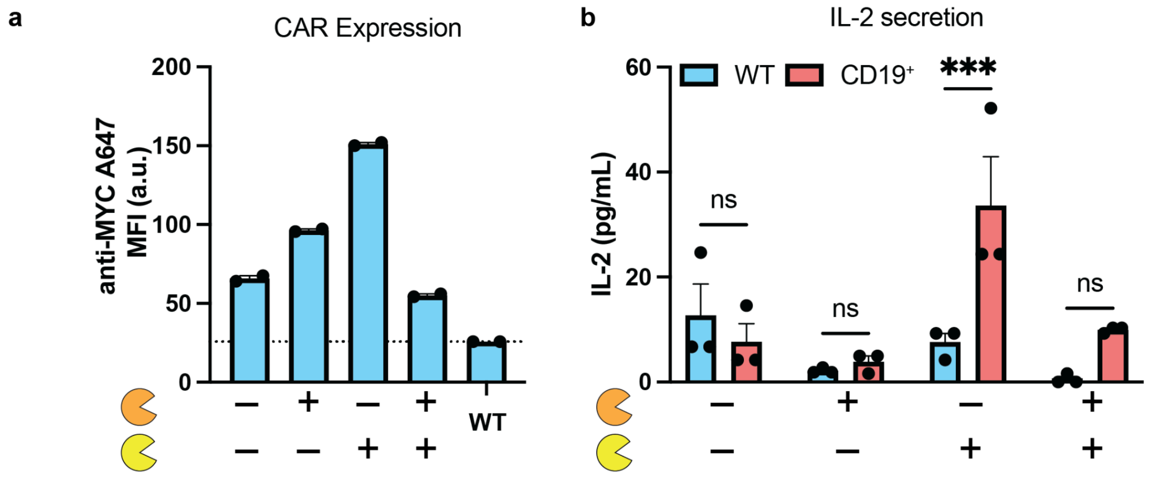


**Supplementary Figure 15:** **a)** Conditional surface expression of CAR validated by surface staining the Myc tag present on the anti-CD19 CAR. n = 2 biological replicates. **b)** Histograms of IL-2 ELISA from supernatant collected from co-culture experiments between engineered Jurkat and K562 cells. IL-2 was significantly increased in the HCVP-only cell line when co-cultured with CD19+ K562 cells. n = 3 biological replicates. Statistical significance was calculated using a two-way ANOVA with Bonferroni's multiple comparisons test. ***$p < 0.001$.

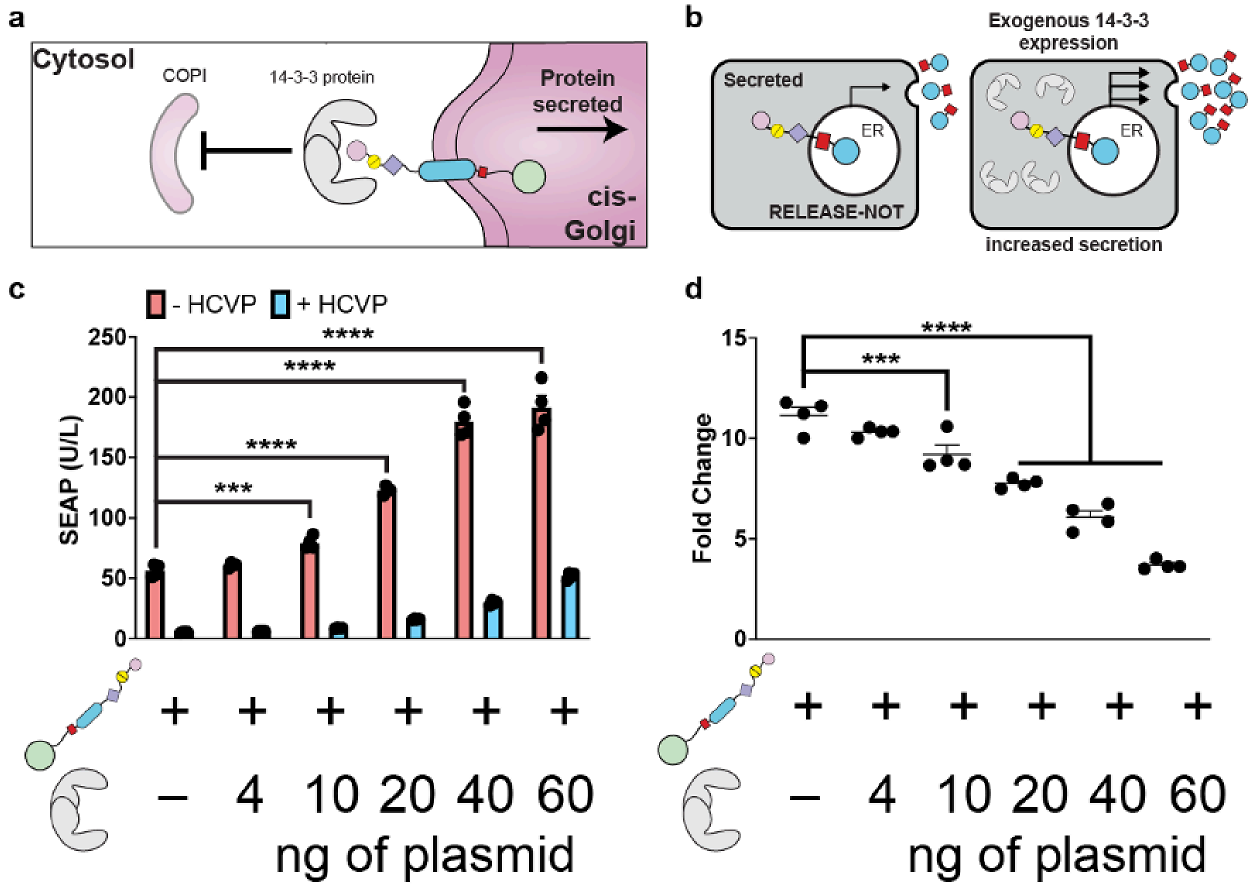

**<u>Supplementary Figure 16:</u>** **a)** Schematic cartoon representing protein secretion controlled via recruitment of 14-3-3 scaffolding proteins by blocking recognition of ER-retention motifs from COPI. **b)** Schematic depicting delivering exogenous 14-3-3 proteins to boost protein expression of RELEASE-NOT. **c)** Dot plots showing SEAP secretion positively correlated with increasing amounts of exogenous 14-3-3ζ co-transfected in HEK293 cells. N = 4 biological replicates. Statistical significance was calculated using a two-way ANOVA with Bonferroni's multiple comparisons test. **d)** The fold change was reduced based on the amount of exogenous 14-3-3ζ co-transfected. N = 4 biological replicates. Statistical significance was calculated using a one-way ANOVA with a Tukey's multiple comparison test. ***$p$ < 0.001, ****$p$ < 0.0001.

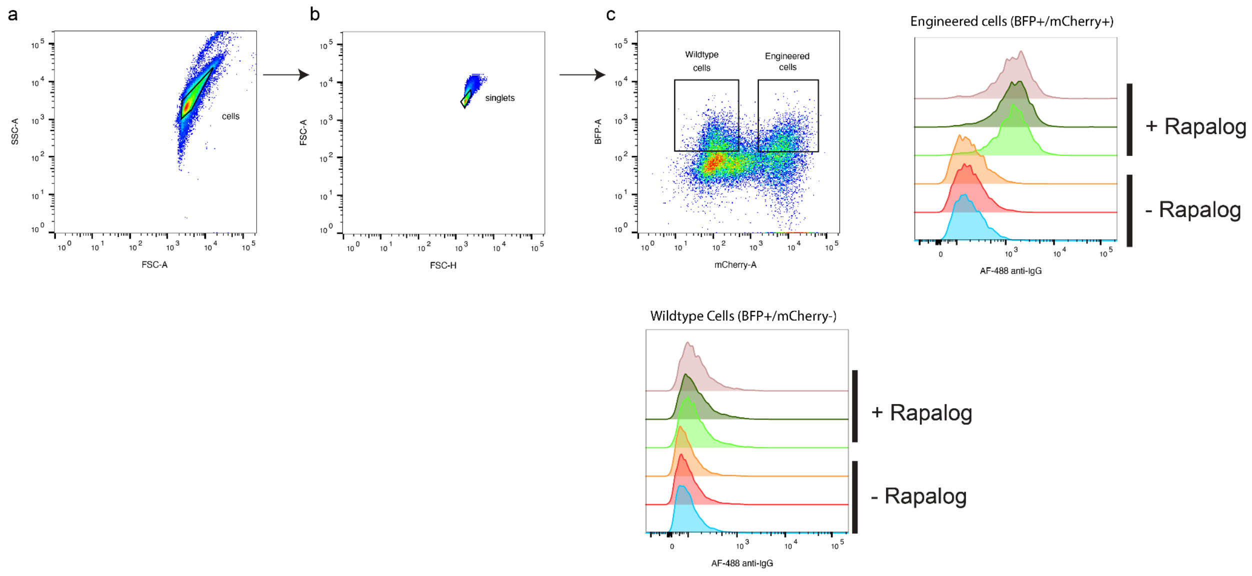


**<u>Supplementary Figure 17:</u>** Representative flow plots to outline gating strategy for experiments with IgG-RELEASE. **a)** Cells were gated based on FSC-A and SSC-A, **b)** followed by gating on singlets. **c)** For experiments with mixed cell populations, engineered cells were gated from wildtype cells based on mCherry expression. We also gated on BFP expression to select for cells that were transfected with IgG-RELEASE-T2A-BFP. Representative experiment analyzing the amount of surface IgG after staining with an anti-IgG AF-488 antibody of engineered (BFP+/mCherry+) and wildtype HEK293 cells (BFP+/mCherry-), incubated with rapalog or vehicle.

### Bibliography

1. Vlahos, A.E., Kang, J., Aldrete, C.A., Zhu, R., Chong, L.S., Elowitz, M.B., and Gao, X.J. (2022). Protease-controlled secretion and display of intercellular signals. Nat. Commun. *13*, 912. 10.1038/s41467-022-28623-y.

**Bandpass and Bandstop Model Derivations**

In order to analyze effects of differential protease cleavage rates on the behaviors of a bandpass architecture, we constructed a simple ordinary differential equation model to evaluate at steady-state. The model incorporates three major types of time-dependent events. First, we consider zeroth order (ie: constant) transcription and translation of the encoded RELEASE protein. We also consider eachs species to have equivalent first-order degradation kinetics. Finally, we model protease cleavage events as bimolecular interactions, equivalent to a Mechalis-Menton model far from saturation. Thus we can approximate our protease cleavage events as:

$$\frac{d[Substrate_{cleav}]}{dt} = \frac{k_{cat}[Protease][Substrate]}{K_m + [Substrate]}$$

In the limit where the substrate concentrations are far from saturating the protease amount, we can assume $[S] << K_m$, and this term can be simplified to:

$$\frac{d[Substrate_{cleave}]}{dt} = \frac{k_{cat}[Protease][Substrate]}{K_m} = k_{cleave}[Protease][Substrate]$$

From this, we can derive a system of equations to describe all possible states our substrate, Bandpass RELEASE, can take. Here, we assume that the RELEASE construct can be in one of four states: 1) cleaved zero times (as translated), 2) cleaved once at the more efficient cutsite, 3) cleaved once at the less efficient cutsite, and 4) cleaved at both cutsite. In this model, we can assume that only the second species, which is cleaved only at the more efficient cleavage site, is secreted out of the cell. We can also assume that retained species 1, 3, and 4, have some amount of leakiness and can still be secreted. We can effectively treat these leaky species as species 2 to simplify the model. We can describe this in the following system of equations:

$$\frac{d[RELEASE_{BP1}]}{dt} = T - [RELEASE_{BP1}]\left(- k_1^{cleave}[TEVp] - \alpha k_1^{cleave}[TEVp] - k_d - k_{leak}\right)$$

$$\frac{d[RELEASE_{BP2}]}{dt} = \left(k_1^{cleave}[TEVp][RELEASE_{BP1}]\right) - [RELEASE_{BP2}]\left(\alpha k_1^{cleave}[TEVp] + k_d\right)$$

$$+ k_{leak}\left([RELEASE_{BP1}] + [RELEASE_{BP3}] + [RELEASE_{BP4}]\right)$$

$$\frac{d[RELEASE_{BP3}]}{dt} = \alpha k_1^{cleave}[TEVp][RELEASE_{BP1}] - [RELEASE_{BP3}]\left( k_1^{cleave}[TEVp] + k_d + k_{leak}\right)$$

$$\frac{d[RELEASE_{BP4}]}{dt} = \alpha k_1^{cleave}[TEVp][RELEASE_{BP1}] + k_1^{cleave}[TEVp][RELEASE_{BP3}] - [RELEASE_{BP4}]\left(k_d + k_{leak}\right)$$

Here, each RELEASE species is denoted as $[RELEASE_{BPn}]$ for bandpass and where $n$ is the species number. $T$ represents constant transcription and translation of an uncleaved

RELEASE-bandpass protein, $k_1^{cleave}$ is the Mechalis-Menton rate constant of protease cleavage far from substrate saturation for the more efficient cleavage site, α is the ratio of the protease cleavage rates between the less:more efficient cleavage site, $k_d$ is the first-order degradation constant assuming all constructs degrade similarly, and $k_{leak}$ is the first order secretion rate of the typically-retained species onto the surface of the cell.

A steady state solution for $[RELEASE_{BP2}]$ can then be found as a function of $[TEVp]$ by solving this system of equations simultaneously, yielding:

$$[RELEASE_{BP2}] = \frac{T(k_d k_{leak} + k_{leak}^2 + k_d k_1^{cleave}[TEVp] + k_{leak} k_1^{cleave}[TEVp] + \alpha k_{leak} k_1^{cleave}[TEVp])}{k_d(k_d + k_{leak} + \alpha k_1^{cleave}[TEVp])(k_d + k_{leak} + k_1^{cleave}[TEVp] + \alpha k_1^{cleave}[TEVp])}$$

By changing the ratio of cleavage efficiencies, α, and setting the various kinetic rate constants, we can probe expected circuit behavior as we alter the cleavage sites.

Similarly, a system of ordinary differential equations can be defined to model the expected interactions in the bandstop circuit. Here, the protein architecture differs slightly, thus we assume the system can be cleaved 1) zero times, 2) once at the more efficient cut site, 3) cut at the less efficient cutsite, and the uncleaved species can also be bound by a scaffold protein. Here we assume the scaffold protein is limiting and binds with first-order Hill kinetics. With these assumption we can define a system of equations:

$$\frac{d[RELEASE_{BS1}]}{dt} = T - \frac{\beta[RELEASE_{BS1}]}{K_d + [RELEASE_{BS1}]} - [RELEASE_{BS1}]\left(-k_1^{cleave}[TEVp] - \alpha k_1^{cleave}[TEVp] - k_d\right)$$

$$\frac{d[RELEASE_{BS1}^{Bound}]}{dt} = \frac{\beta[RELEASE_{BS1}]}{K_d + [RELEASE_{BS1}]} - [RELEASE_{BS1}^{Bound}]\left(-k_1^{cleave}[TEVp] - \alpha k_1^{cleave}[TEVp] - k_d\right)$$

$$\frac{d[RELEASE_{BS2}]}{dt} = k_1^{cleave}[TEVp]\left([RELEASE_{BS1}] + [RELEASE_{BS1}^{Bound}]\right) - [RELEASE_{BS2}]\left(\alpha k_1^{cleave}[TEVp] + k_d\right)$$

$$\frac{d[RELEASE_{BS3}]}{dt} = \alpha k_1^{cleave}[TEVp]\left([RELEASE_{BS1}] + [RELEASE_{BS1}^{Bound}] + [RELEASE_{BS2}]\right) - k_d[RELEASE_{BS3}]$$

Here, again each RELEASE species is denoted as $[RELEASE_{BSn}]$ for bandstop and where $n$ is the species number, and species 1 can exist either bound or unbound, as noted. Again, $T, k_1, k_d$, and α are defined as before. Additionally, β and $K_d$ are the Hill binding maximum and equilibrium dissociation constants for the scaffolding protein binding to the uncleaved $[RELEASE_{BS1}]$ species. From this, we can use a computational solver to determine the steady state solution of this system as a function of $[TEVp]$ and analyze the sum of the two secreting species, $[RELEASE_{BS1}^{Bound}]$ and $[RELEASE_{BS3}]$ as our total expected output at a given amount of protease.
